# Thirty days of supplementation with PQQ reprograms immunometabolic networks in Western diet-fed female baboons

**DOI:** 10.64898/2026.02.12.705540

**Authors:** Sunam G. Dockins, Kimberly E. Hyatt, Darlene N. Reuter, Taylor L. Stevens, Amanda B. Tresler, James F. Papin, Dean A. Myers, Karen R. Jonscher

## Abstract

Western-style diets promote chronic metabolic inflammation and dyslipidemia, yet safe interventions that restore immunometabolic homeostasis remain limited. Pyrroloquinoline quinone (PQQ) is a naturally occurring redox cofactor with antioxidant and metabolic regulatory properties, but its systemic effects in translational preclinical models are poorly defined. Here, we examined the impact of short-term PQQ supplementation in obese adult female olive baboons (*Papio anubis*) chronically fed a Western diet. Using a human-equivalent dose administered for 30 days, we found that PQQ supplementation significantly reduced circulating markers of systemic inflammation and cholesterol in Western-diet-fed animals, lowering circulating C-reactive protein, soluble CD163, and atherogenic lipoprotein fractions independent of changes in adiposity. Proteomic and pathway analyses of circulating proteins in plasma and serum revealed suppression of complement, thrombo-inflammatory, and extracellular matrix remodeling pathways, alongside enhanced lipoprotein assembly, remodeling, and clearance. Network analyses identified restoration of neurotrophic tyrosine kinase receptor 1 (NTRK1)- and forkhead box A2 (FOXA2)-regulated signaling as central features of the PQQ response, accompanied by inhibition of pro-fibrotic, xenobiotic, and inflammatory pathways, as well as predicted activation of liver X receptor (LXR)- and insulin growth factor (IGF)-associated metabolic programs. These findings demonstrate that PQQ rapidly reprograms systemic immunometabolic networks in a nonhuman primate model of diet-induced metabolic stress, highlighting FOXA2- and neurotrophin-associated pathways as novel targets of PQQ’s action.

## INTRODUCTION

The increasing prevalence of obesity and its associated complications represents a major global public health challenge. In the United States (US), approximately 42% of adults meet criteria for obesity ^1^. Pediatric and maternal obesity rates are also rising rapidly. According to the Centers for Disease Control and Prevention (CDC) childhood obesity facts, nearly 14.7 million children and adolescents between the ages of 2-19 years have obesity, which in children is defined as having a body mass index (BMI) at or above the 95^th^ percentile for age and sex ^2^. Among women of reproductive age, almost 30% enter pregnancy with obesity, reflecting an ∼11% increase across maternal ages, educational backgrounds, and races compared with estimates prior to 2016 ^3^. Maternal obesity is strongly associated with adverse outcomes in offspring, including gestational diabetes, pre-eclampsia, pre-term birth, small and large for gestational age, and infant mortality ^4–6^. If current trends persist, projections suggest that by 2030 roughly half of the adult population in the US will be living with overweight or obesity ^7^.

The chronic nature of overweight and obesity, together with their many comorbidities, generates substantial health-care expenditures and imposes significant economic burdens on patients, health systems, and public health infrastructure ^8,9^. While lifestyle modification, bariatric surgery, and newer glucagon-like peptide-1 receptor agonists (GLP-1Ras) can be effective for some individuals, these approaches carry risk, are not universally accessible, and may be unsuitable for broad implementation ^10^. Importantly, GLP-1 RAs are currently contraindicated for pregnancy and lactation, leaving few viable therapeutic options for maternal and pediatric obesity. The development of safe, affordable, and scalable interventions therefore remains a critical priority in addressing this escalating global health threat.

Pyrroloquinoline quinone (PQQ) has emerged as a bioactive compound with beneficial effects on metabolic health in obesity ^11–13^. Originally identified in the 1960s as a redox cofactor for bacterial dehydrogenases ^14^, PQQ is present in a variety of foods and is considered an essential nutritive factor with vitamin-like biochemical properties ^15^. Studies have shown PQQ to have potent antioxidant and anti-inflammatory activity ^15–18^, targeting essential biological processes, including mitochondrial biogenesis ^16–18^, lipid and glucose metabolism ^19^, reproduction ^20,21^, and growth and aging ^22–25^. Our group has previously shown beneficial effects of PQQ in a mouse model of Western diet (WD)-induced obesity, where both pre- and post-natal PQQ administration to dams protected offspring from WD-induced hepatic oxidative stress, inflammation, and lipotoxicity, conferring durable resistance to hepatic steatosis and metabolic dysfunction-associated steatotic liver disease (MASLD) ^26^.

The objective of the present study was to evaluate the antioxidant, anti-inflammatory, and metabolic effects of PQQ in a non-human primate (NHP) model, the Olive baboon (*Papio anubis*). Non-human primates are a highly translational model for investigating complex polygenetic metabolic diseases such as obesity because of their genetic, epigenetic, and developmental similarities to humans ^27,28^. In this study, we administered PQQ daily for 30 days to non-pregnant adult female olive baboons fed either a control diet (CD, n=5) or Western-style diet (WD, n=4) to evaluate its efficacy in improving systemic metabolic and inflammatory health. We quantified systemic metabolic (triglycerides and cholesterol) and inflammatory (CRP, sCD63) markers by ELISA and performed proteomic profiling in serum and plasma collected at baseline and after 30 days of PQQ supplementation. Our results show that 30 days of supplementation with a PQQ dose that is routinely suggested for daily human use (0.25 mg/kg body weight) led to improved metabolic and inflammatory profiles in WD-fed obese baboons.

## RESULTS

PQQ was administered to non-pregnant female baboons daily, packaged into a cakeball treat (**Figure 1a, 1b**). Plasma and serum were collected at D0 (baseline) and after 30 days of supplementation.

**Figure 1.**
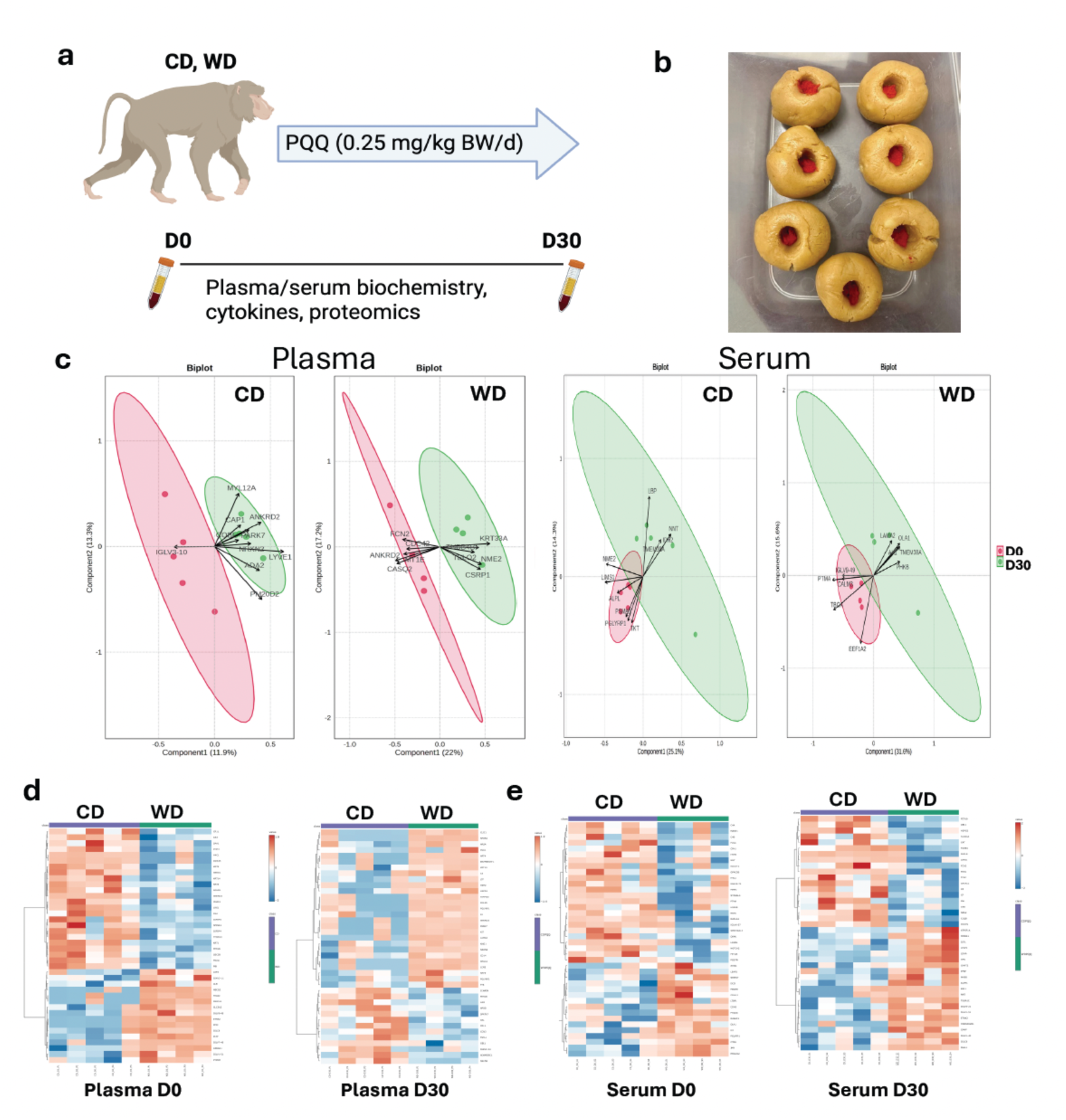
Thirty days of PQQ supplementation alters the distribution of circulating proteins in plasma and serum. a) Non-pregnant female baboons fed either CD or WD for 18 months were supplemented with 0.25 mg/kg/body weight/day of PQQ. Plasma and serum were collected at baseline and on day 30. b) Cake balls used to deliver PQQ to baboons. c) Paired partial least squares discriminant analysis of circulating peptides in plasma and serum, comparing between day 0 and day 30 for each diet group. Heatmaps in d) plasma and e) serum obtained using unpaired analysis of circulating proteins. Unpaired analysis on day 0 and day 30 compared between diet groups. n=4-5/group.

### Anthropometrics

There were no differences in body weights between the CD and WD female baboons at the start or end of the study. There is considerable variability in body weights and lengths in adult female (and male) *Papio anubis* in our cohort, precluding the use of weight alone as a proxy for obesity. There was a significant difference in the sum of skin folds (SSF) between CD and WD cohorts at the study’s start and end (**Table 1)** reflecting a 34% increase in subcutaneous adiposity between dams fed chronic WD compared with CD. PQQ supplementation for 30 days did not affect body weight or SSF in either group. Sum of skin folds did not change in either diet group between study start and finish.

**Table 1.**
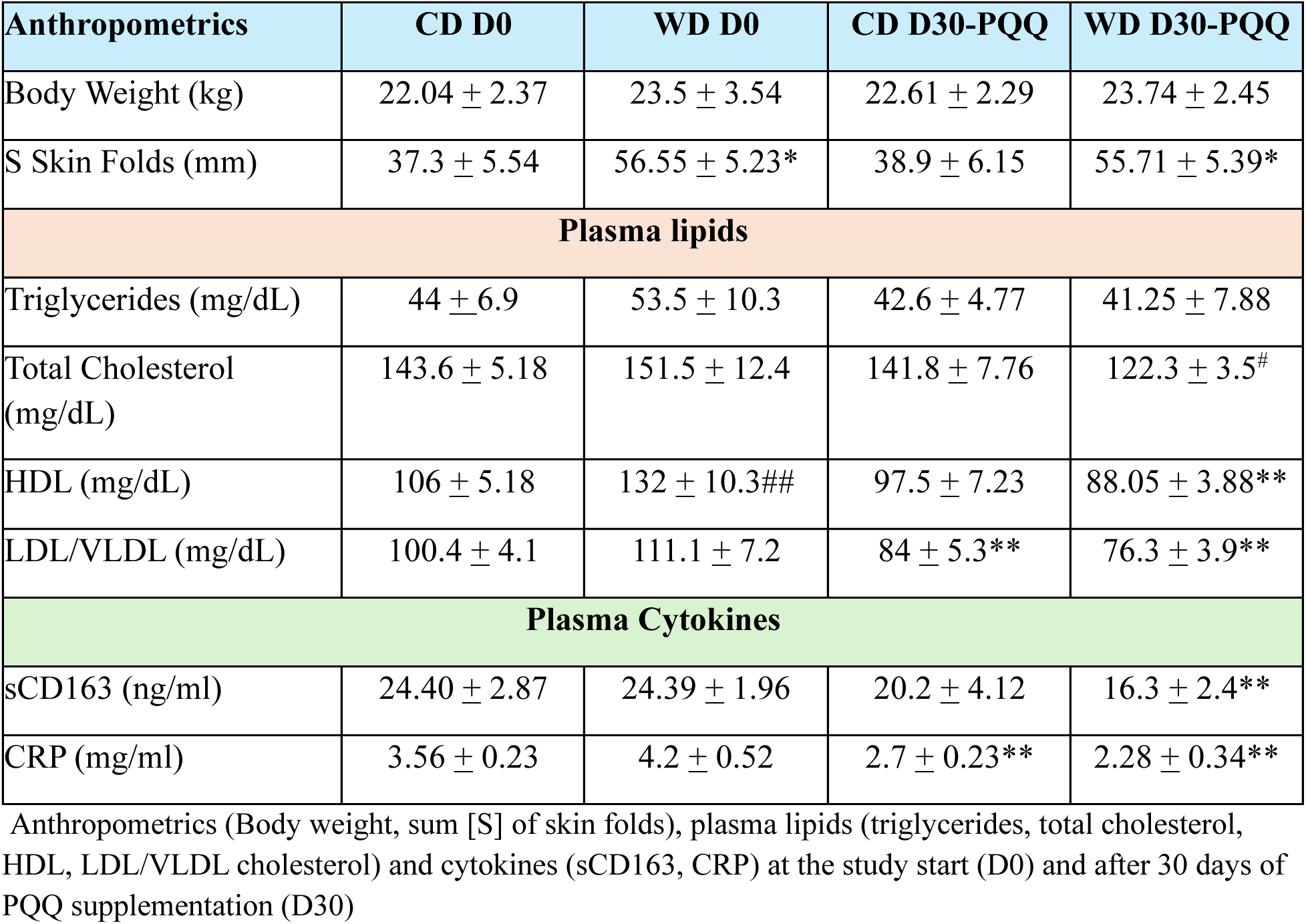
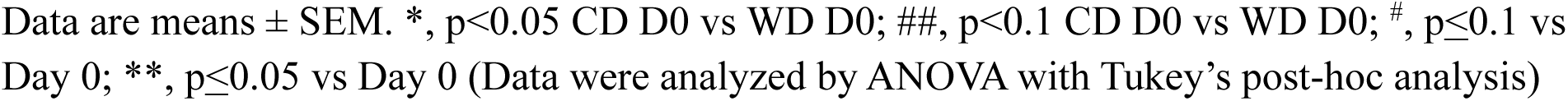
Baboon anthropometrics and plasma lipids and cytokines.

| Anthropometrics | CD D0 | WD D0 | CD D30-PQQ | WD D30-PQQ |
| --- | --- | --- | --- | --- |
| Body Weight (kg) | 22.04 ± 2.37 | 23.5 ± 3.54 | 22.61 ± 2.29 | 23.74 ± 2.45 |
| S Skin Folds (mm) | 37.3 ± 5.54 | 56.55 ± 5.23* | 38.9 ± 6.15 | 55.71 ± 5.39* |
| **Plasma lipids** |  |  |  |  |
| Triglycerides (mg/dL) | 44 ± 6.9 | 53.5 ± 10.3 | 42.6 ± 4.77 | 41.25 ± 7.88 |
| Total Cholesterol (mg/dL) | 143.6 ± 5.18 | 151.5 ± 12.4 | 141.8 ± 7.76 | 122.3 ± 3.5<sup>#</sup> |
| HDL (mg/dL) | 106 ± 5.18 | 132 ± 10.3<sup>##</sup> | 97.5 ± 7.23 | 88.05 ± 3.88<sup>**</sup> |
| LDL/VLDL (mg/dL) | 100.4 ± 4.1 | 111.1 ± 7.2 | 84 ± 5.3<sup>**</sup> | 76.3 ± 3.9<sup>**</sup> |
| **Plasma Cytokines** |  |  |  |  |
| sCD163 (ng/ml) | 24.40 ± 2.87 | 24.39 ± 1.96 | 20.2 ± 4.12 | 16.3 ± 2.4<sup>**</sup> |
| CRP (mg/ml) | 3.56 ± 0.23 | 4.2 ± 0.52 | 2.7 ± 0.23<sup>**</sup> | 2.28 ± 0.34<sup>**</sup> |

Anthropometrics (Body weight, sum [S] of skin folds), plasma lipids (triglycerides, total cholesterol, HDL, LDL/VLDL cholesterol) and cytokines (sCD163, CRP) at the study start (D0) and after 30 days of PQQ supplementation (D30)
Data are means ± SEM. \*, $p < 0.05$ CD D0 vs WD D0; ##, $p < 0.1$ CD D0 vs WD D0; #, $p \leq 0.1$ vs Day 0; \*\*, $p \leq 0.05$ vs Day 0 (Data were analyzed by ANOVA with Tukey's post-hoc analysis)

### Serum lipids

Due to variability in the WD-fed females, there were no significant differences between the two dietary groups in serum triglycerides, total cholesterol, or LDL/VLDL cholesterol at Day 0; however, HDL cholesterol was significantly elevated (p < 0.05) in the WD females compared with CD at Day 0. PQQ supplementation for 30 days reduced serum triglycerides by 3% in CD and 23% in WD, although differences were not significant. PQQ supplementation significantly reduced total cholesterol (p=0.024), HDL (p=0.0009) and LDL/VLDL (p=0.0005) cholesterol in the WD females (33% and 31% vs. Day 0, respectively). In contrast, the effect of PQQ on HDL and VLDL cholesterol levels in CD females was a very modest reduction (8% and 16% compared with Day 0) that was only significant for LDL/VLDL cholesterol (p=0.034).

### Serum cytokines

There were no significant differences between diet groups in serum CRP or sCD163 (a marker for monocyte/macrophage activation) on Day 0. PQQ supplementation for 30 days significantly lowered both CRP (p=0.043) and sCD163 (p=0.045) in WD (46% and 33% compared with Day 0) but only significantly lowered CRP (24%, p=0.031), and not sCD163 (17%) in CD.

### Proteomics

To investigate the effect of PQQ on the circulating proteome, untargeted proteomics was performed using LC-MS/MS in plasma and serum collected at Day 0 and Day 30 of the study. Up to 732 proteins were identified in serum and 705 proteins in plasma.

#### Multivariate statistical analysis

We first used the Statistical Analysis [one factor] module in MetaboAnalyst 6.0 to perform a paired analysis comparing plasma with serum, using all groups, to determine whether differences in the protein profiles between plasma and serum were diet- or PQQ-related (**Supplementary Fig. 1a**). Partial Least Squares Discriminant Analysis (PLS-DA) showed that features in serum and plasma samples clustered separately, with further separation based on PQQ for CD in plasma but not serum. Paired comparisons between D0 and D30 did not reveal further clustering by diet or sample origin (**Supplementary Fig. 1b**). Unpaired PLS-DA analysis showed that features clustered by diet in WD v CD in both plasma and serum (**Supplementary Fig. 1c, 1d**). We next used pairwise comparisons in serum and plasma separately to identify proteins promoting the clustering. PLS-DA biplots show that markedly different proteins drive the separation between D0 and D30 in plasma and serum for each diet group (**Figure 1c**).

Using relaxed stringency for significance, volcano plots in both unpaired (WD D0 v CD D0) and paired (CD D30 v CD D0, WD D30 v WD D0) comparisons showed unique protein changes in plasma (**Supplementary Table 2**) and serum (**Supplementary Table 3**). In plasma, expression of ankyrin repeat domain 2 (ANKRD2) was significantly reduced by WD (p=0.01) and markedly elevated by PQQ (p=0.053). No other proteins were significantly changing in more than one pairwise comparison in plasma or serum.

Unpaired hierarchical clustering of the top 40 proteins (determined by t-test p value) was used to compare diets in plasma and serum at D0 and D30 (**Figure 1d, 1e**). Clustering based on diets was observed at both D0 and D30; however, the list of top 40 proteins was different between groups. In plasma, the effect of diet was shown in the relative abundances of Complement C3b/C4b Receptor 1 Like (CR1L), Fetuin B (FETUB), and Scavenger Receptor Cysteine Rich Family Member With 5 Domains (SSC5D), which were reduced in WD compared to CD, regardless of PQQ. Relative abundance of ANKRD2 was increased in WD compared with CD at D30, consistent with the volcano plot analysis. Relative abundance of Keratin 9 (KRT9) was also increased by PQQ. In serum, FETUB and Periostin (POSTN) were decreased in WD compared with CD at both D0 and D30. Nicotinamide Nucleotide Transhydrogenase (NNT) showed an effect of PQQ in serum, with relative abundance in WD compared with CD decreased at D0 and increased at D30. Five proteins were common among the top 40 between comparisons in serum (FETUB, C4B, POSTN, LEAP2, AHSG). However, their patterns of expression in comparison between WD and CD at D0 and D30 did not change.

Paired analyses in plasma showed that Carbonic anhydrase (CA1, CA2) and Hemoglobin (HBA, HBB) were increased at D30 in CD, with more variable results in WD (**Supplementary Fig. 2a**). In serum, IGHV3-48, IGKV3-11 and EFEMP1 (encoding Fibulin-3) were decreased by PQQ in both CD and WD, while an effect of diet was observed for Orosomucoid-1 (ORM1) (**Supplementary Fig. 2b**).

Together, these results suggest that dietary state and PQQ supplementation produce distinct molecular responses depending on the physiological context (diet, treatment, plasma vs. serum). This suggests that PQQ’s effects, and diet-induced metabolic changes, are tissue- and pathway-specific, rather than global.

#### Functional Annotation and Enrichment

We next sought to identify enriched functional categories and predicted protein-protein interactions associated with PQQ supplementation. Protein IDs were mapped to gene symbols using UniProt ID Converter or BLAST. Features with a negative value of log2 fold change between comparisons (downregulated) and those with a positive value of log2 fold change (upregulated) were analyzed separately using DAVID and STRING (**Figure 2a**). The number of mapped and unmapped IDs for DAVID and STRING analyses was summarized in **Supplementary Table 1**, as were the number of clusters and the number of unclustered terms for the DAVID analyses.

**Figure 2.**
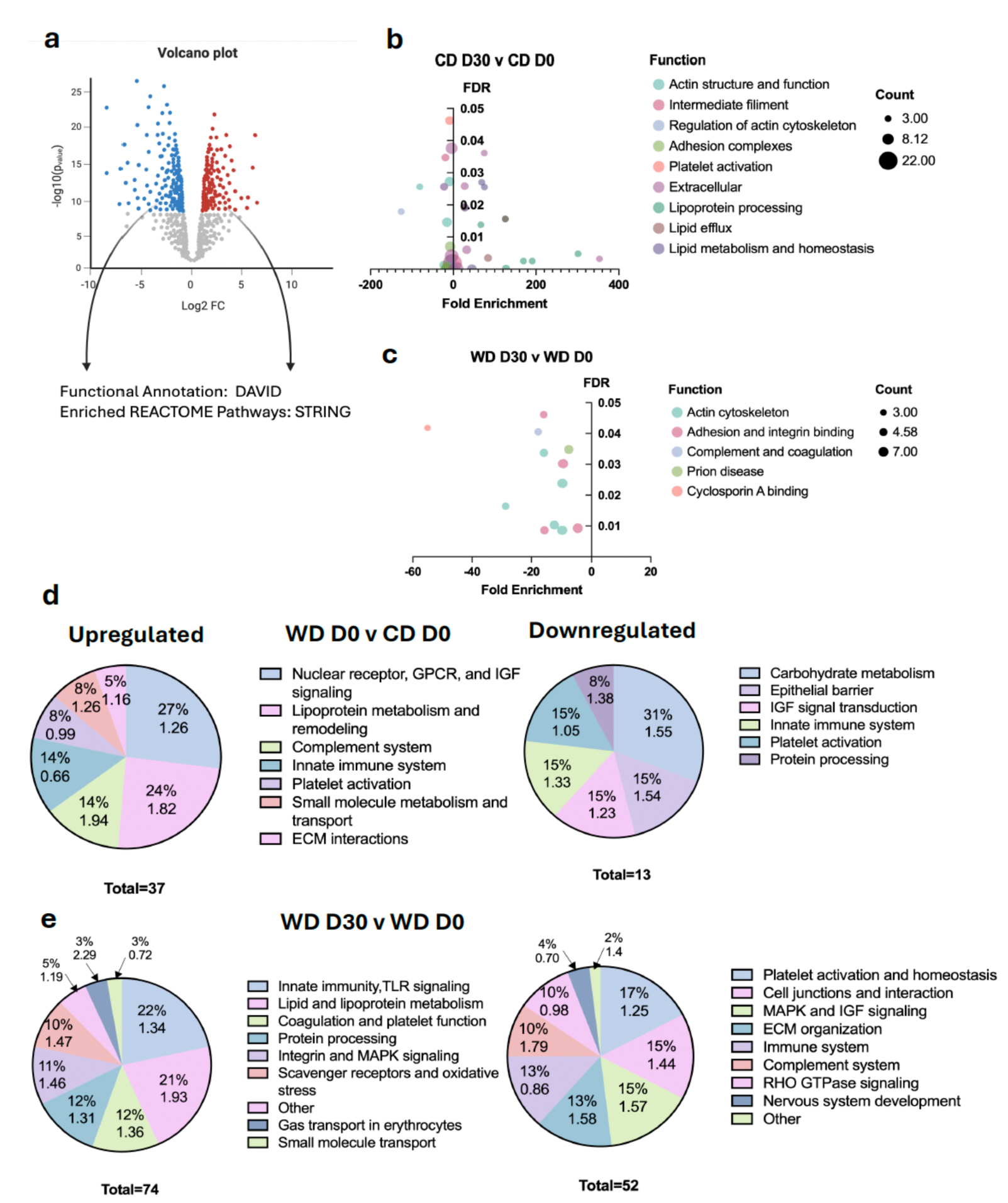
Functional annotations of up- and down-regulated proteins using DAVID and STRING reveal a PQQ-mediated effect on nuclear receptor signaling, lipid metabolism, and the innate immune system. a) Up- and down-regulated proteins (p<0.1, log2 fold-change > |1|) were separately analyzed using DAVID and STRING for each diet group combination. Significantly enriched functions from DAVID analysis (FDR<0.05) in b) CD and c) WD, comparing between day 0 and day 30. Significantly enriched Reactome pathways associated with up- and downregulated proteins in d) WD compared with CD at day 0 and e) WD at day 30 following PQQ supplementation compared with day 0.

Analysis with DAVID showed several significantly enriched functional annotation terms in plasma and serum for each pairwise comparison (**Figure 2b, 2c; Supplementary Table 4)**. In CD plasma (**Figure 2b**), the fold enrichment of pathways involved in cholesterol transport and lipid clearance was highly increased by PQQ, whereas fold enrichment of actin cytoskeleton remodeling and cell adhesion terms, including focal adhesion, adherence junction, and platelet activation, were modestly diminished. In WD plasma, proteins associated with annotations involved in integrin binding, actin remodeling, and focal adhesion signaling were also attenuated by PQQ, and in WD serum, proteins involved in the complement and coagulation cascades were downregulated (**Figure 2c**). These data indicate that PQQ may act to attenuate systemic inflammation induced by WD through targeting cellular adhesion.

Enriched Reactome pathways were obtained from the STRING analysis of up and downregulated proteins from each comparison in plasma and serum (**Figure 2d, 2e**; **Supplementary Table 5)**. Over 50% of the enriched Reactome pathways predicted to be upregulated by WD at D0, compared with CD, were associated with nuclear receptor, G-protein coupled receptor, and insulin growth factor (IGF) signaling, in part through the xenobiotic transcription factor Liver X Receptor (LXR; **Figure 2d**). HDL cholesterol remodeling and transport were also highly enriched, whereas pathways predicted to be downregulated primarily included those associated with carbohydrate metabolism. Pathways associated with the epithelial barrier, the innate immune system, and platelet activation were predicted to be downregulated. Almost half of the enriched Reactome pathways predicted to be upregulated by PQQ in WD-fed baboons were involved in innate immunity, TLR signaling, and lipid and lipoprotein metabolism (**Figure 2e**). Predicted downregulated pathways involved in thrombo-inflammatory signaling were attenuated by lowering integrin-mediated adhesion, platelet activation and degranulation, and complement activity, in alignment with findings from the DAVID analysis.

#### Pathway Analysis

Because DAVID and STRING identify enriched pathways and protein–interaction networks but do not provide directional or mechanistic insight, we performed Ingenuity Pathway Analysis (IPA) to predict upstream regulatory drivers, assess canonical pathway activation states, and evaluate potential physiological consequences of the observed protein changes. Significance stringency was relaxed to allow for analysis of a minimum of 100 molecules. The number of mapped analysis-ready proteins that were up- and down-regulated, and the p value cutoffs are summarized in **Supplementary Table 1**. Volcano plots show up- and down-regulated proteins with |log2 fold change| > 1 comparing WD v CD at baseline in plasma (**Figure 3a**) and serum (**Figure 3c**), and the effect of PQQ in CD- (**Supplementary Fig. 3a, 3b**) and WD-fed baboons (**Figure 3b, 3d**).

**Figure 3.**
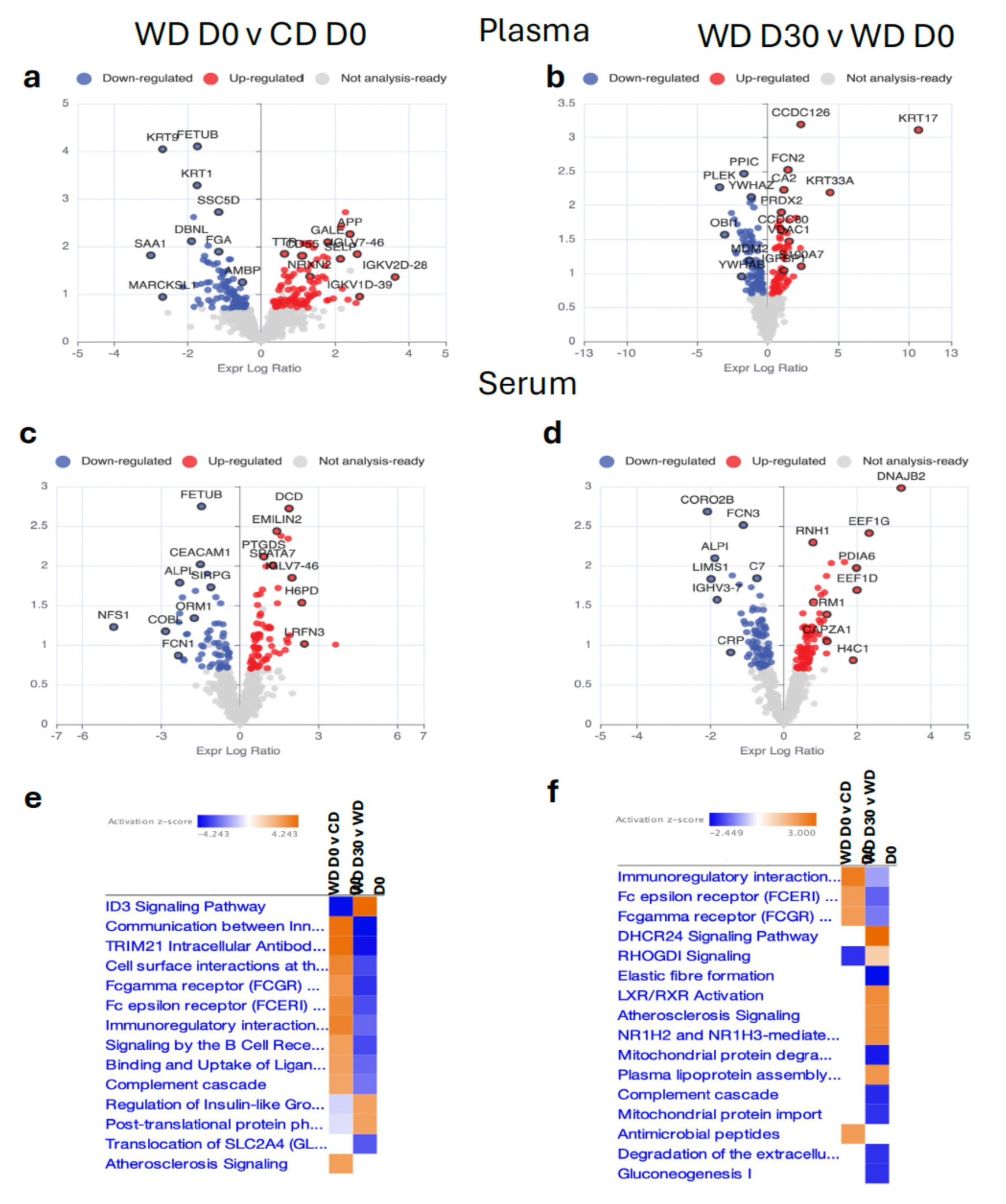
Protein composition and associated canonical pathways inferred using Ingenuity Pathway Analysis (IPA) are markedly different between plasma and serum. Volcano plots were generated for comparisons between diet groups in plasma at a) baseline (WD D0 v CD D0) and b) between baseline and PQQ supplementation in WD (WD D30 v WD D0). Similar comparisons were made in serum (c, d). Enriched canonical pathways were identified in e) plasma and f) serum.

Canonical pathway analysis was performed in plasma and serum using both “Core” and “Comparison” analyses functions in IPA (**Supplementary Table 6, 7**; **Figure 3e, 3f**). In plasma, a total of 74 enriched canonical pathways (-log10p > 1.3, |z-score| > 1) were identified between WD D0 v CD D0, CD D30 v CD D0, and WD D30 v WD D0 comparisons (**Supplementary Table 6**). There were 29 pathways that were enriched comparing WD D30 with WD D0, and 20 pathways enriching comparing CD D30 with CD D0. Pathways that were enriched at Day 30, regardless of diet, included ‘*Translocation of SLC2A4 (GLUT4) to the plasma membrane*’, ‘*Intrinsic Pathway for Apoptosis*’, ‘*L1CAM interactions*’, ‘*Integrin Signaling*’, ‘*RHOA Signaling*’, ‘*TP53 Regulated Metabolic Genes*’, ‘*EPH-Ephrin signaling*’, ‘*Smooth Muscle Contraction*’, and ‘*Regulation of Insulin-like Growth Factor (IGF) transport and uptake by IGFBPs*’. These findings suggest PQQ may enhance insulin signaling, glucose utilization, and cell-matrix communication, regardless of diet.

Seventeen pathways were significantly enriched in the comparison between both WD and CD at D0 and between WD D30 and WD D0 (**Table 2**); in 14 of these pathways the z-score was reversed by PQQ, i.e. PQQ treatment reversed or blunted the effect of WD feeding in obese dams. Enriched pathways with |z-score| > 2 that were reversed by PQQ include ‘*Binding and Uptake of Ligands by Scavenger Receptors*’, ‘*Communication between Innate and Adaptive Immune Cells*’, ‘*Complement cascade*’, ‘*Fc epsilon receptor (FCERI) signaling*’, ‘*Fcgamma receptor (FCGR) dependent phagocytosis*’, ‘*ID3 Signaling Pathway*,’ ‘*Immunoregulatory interactions between a Lymphoid and a non-Lymphoid cell*’, ‘*Mitochondrial protein import*’, ‘*Sheddase Signaling Pathway*’, ‘*Signaling by the B Cell Receptor (BCR)*’, and ‘*TRIM21 Intracellular Antibody Signaling Pathway*’. Similar results were found when comparing WD D0 v CD D0 and WD D30 v CD D30 (**Supplementary Fig. 3c**). These data suggest that PQQ reduces WD-driven increase in innate immune activation and inflammation amplified by complement.

**Table 2.** Enriched canonical pathways in plasma and serum.

| Ingenuity Canonical Pathways | -log(p-value) | z-score | -log(p-value) | z-score |
| --- | --- | --- | --- | --- |
| ABRA Signaling Pathway | 3.49 | -1.342 | 3.67 | -1.342 |
| Actin Cytoskeleton Signaling | 2.88 | -1.342 | 7.44 | -1.667 |
| **Antimicrobial peptides** | 6.03 | -1.342 | 6.26 | 1.633 |
| **Binding and Uptake of Ligands by Scavenger Receptors** | 22.2 | 2.683 | 8.64 | -2.53 |
| **Cell surface interactions at the vascular wall** | 15.3 | 3.441 | 9.08 | -3.051 |
| **Communication between Innate and Adaptive Immune Cells** | 5.22 | 4.025 | 4.68 | -4.243 |
| **Complement cascade** | 23.5 | 2.558 | 10.3 | -2.309 |
| **Fc epsilon receptor (FCERI) signaling** | 11 | 3.357 | 7.19 | -3 |
| **Fc gamma receptor (FCGR) dependent phagocytosis** | 13.6 | 3 | 9.35 | -3.464 |
| **ID3 Signaling Pathway** | 5.61 | -4.146 | 4.52 | 4.243 |
| **Immunoregulatory interactions between a Lymphoid and a non-Lymphoid cell** | 11.8 | 3.5 | 6.03 | -2.53 |
| **Mitochondrial protein degradation** | 4.27 | -1.633 | 2.5 | 2 |
| **Mitochondrial protein import** | 3.01 | -2 | 4.3 | 2.236 |
| Response to elevated platelet cytosolic Ca<sup>2+</sup> | 13.8 | -1.069 | 15.8 | -2 |
| **Sheddase Signaling Pathway** | 2.58 | 1.633 | 2.78 | -2.449 |
| **Signaling by the B Cell Receptor (BCR)** | 7.64 | 2.714 | 10.3 | -3.051 |
| **TRIM21 Intracellular Antibody Signaling Pathway** | 8.74 | 4.025 | 9.44 | -4.025 |

**Bold text** indicates pathways with z-score reversed by PQQ

Analysis of canonical pathways in serum showed no overlap of enriched pathways between WD v CD at Day 0 and WD D30 v WD D0 (**Supplementary Table 7**). Thirty-one enriched pathways were identified in total in CD D30 v CD D0 and WD D30 v WD D0. Five pathways were common between both comparisons, four of which were also identified in plasma. ‘*Degradation of the extracellular matrix*’ was uniquely identified as downregulated in serum. Comparison analysis of WD D0 v CD D0 and WD D30 v CD D30 (**Supplementary Fig. 3e**) also showed pathways were similarly enriched, regardless of PQQ, except for ‘*ID3 Signaling Pathway*’ and ‘*Detoxification of Reactive Oxygen Spe*cies’, which were significantly decreased following PQQ treatment.

#### Upstream regulator analysis

To gain more insight into biological mechanisms that may underline the proteomics results, we performed Upstream Regulator analyses in both plasma and serum. Using IPA’s AI mode from the ‘Core Analysis’ module, bubble charts were generated in plasma (**Figure 4a, 4b**) and serum (**Figure 5a, 5b**) using a −log10 p-value cutoff of 2.5 and |z-score| cutoff of 2.0. Interacting networks of predicted upstream regulator interactions were plotted for several of the putative upstream regulators (**Supplementary Fig. 4, 5**). WD feeding led to an enrichment in pathways in plasma activated by TGFBR2, SKIC2, NR1H4 (FXR), and SIX1. Significantly enriched regulators of inhibited pathways included SMARCA4, INSR, STK11 (LKB1), and PPARGC1A. Upstream regulators of pathways activated by PQQ treatment in WD females included PTEN, TXNIP, FOXA2, CST5, and CARM1. Sixteen upstream regulators with |z-score| > 1 were inhibited by PQQ (**Table 3**), consistent with a broad PQQ-mediated suppression of inflammatory and fibrotic signaling networks. Bubble charts based on molecular type showed WD most significantly increased enzymes and ligand-dependent nuclear receptors, and decreased kinases transcription regulators, cytokines (**Figure 4a**). In contrast, PQQ significantly decreased transcription regulators in WD-fed animals (**Figure 4b**).

**Figure 4.**
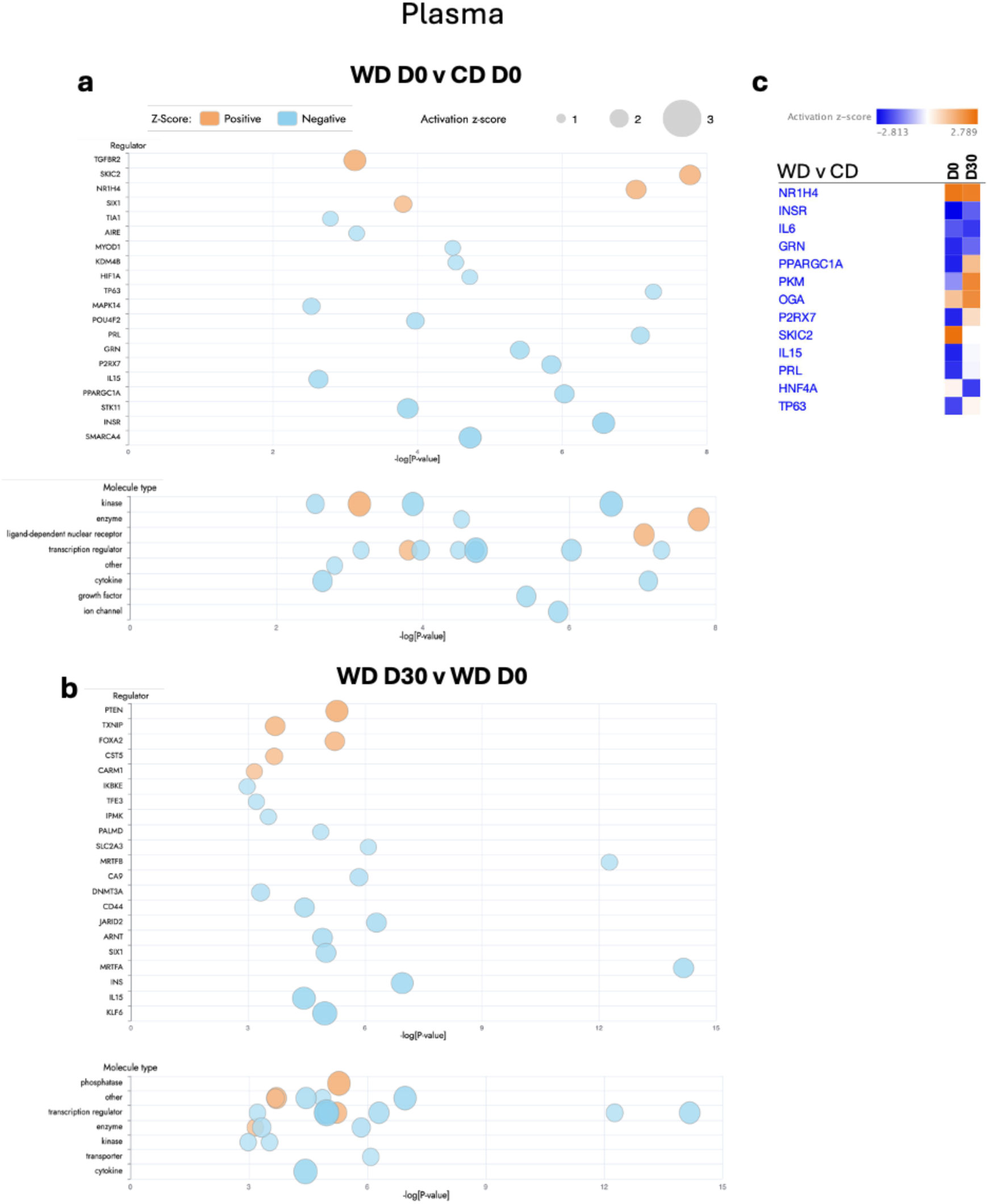
Upstream regulator analysis in plasma. Upstream regulator analysis using IPA comparing between a) diet groups at baseline and b) between baseline and PQQ supplementation in WD. Regulators and their molecule types predicted to be activated (orange) or inhibited (blue) are shown in bubble plots. c) Upstream regulators compared between diets at baseline (D0) and after PQQ supplementation (D30).

**Figure 5.**
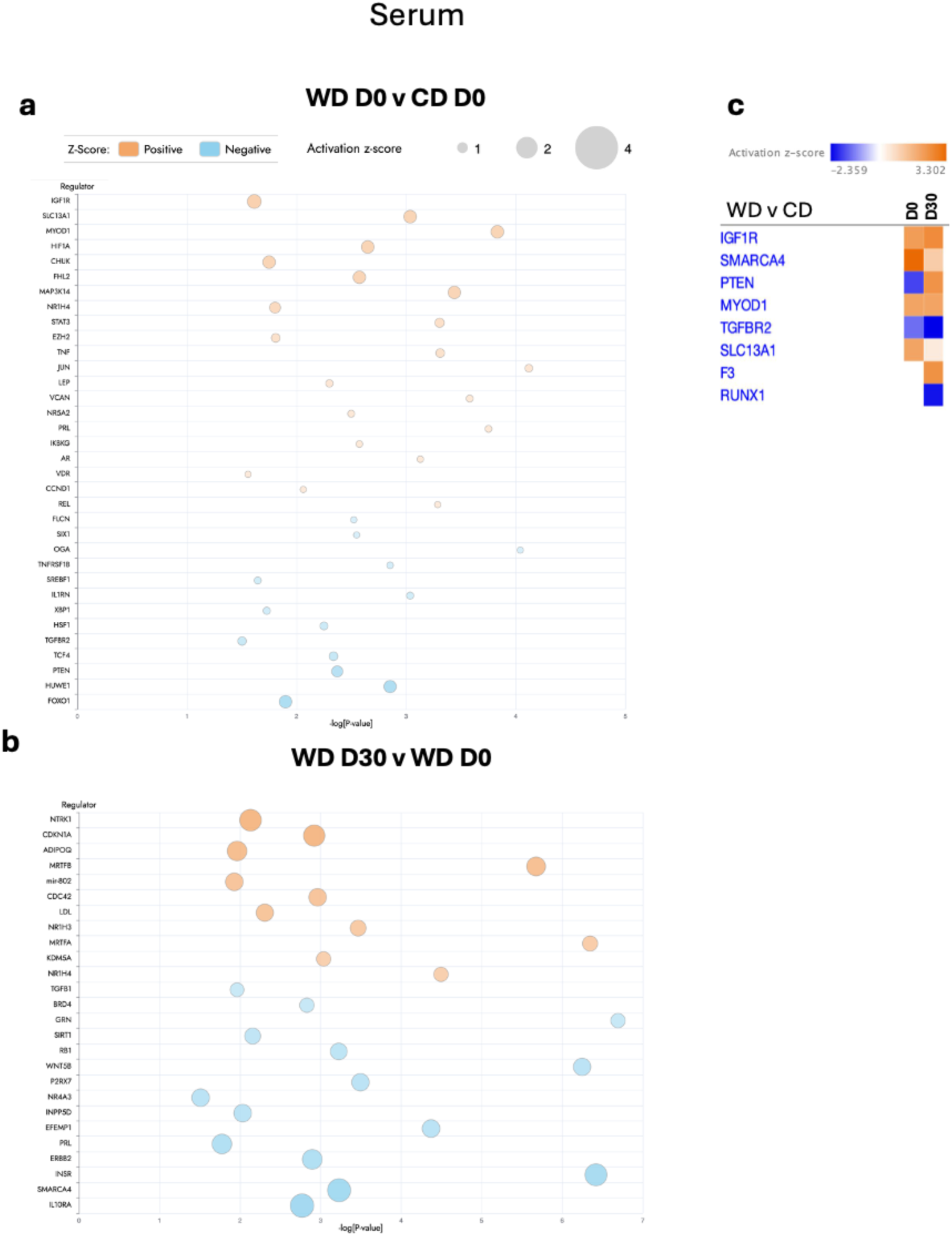
Upstream regulator analysis in serum. Upstream regulator analysis using IPA comparing between a) diet groups at baseline and b) between baseline and PQQ supplementation in WD. Regulators predicted to be activated (orange) or inhibited (blue) are shown in bubble plots. c) Upstream regulators compared between diets at baseline (D0) and after PQQ supplementation (D30).

**Table 3.**
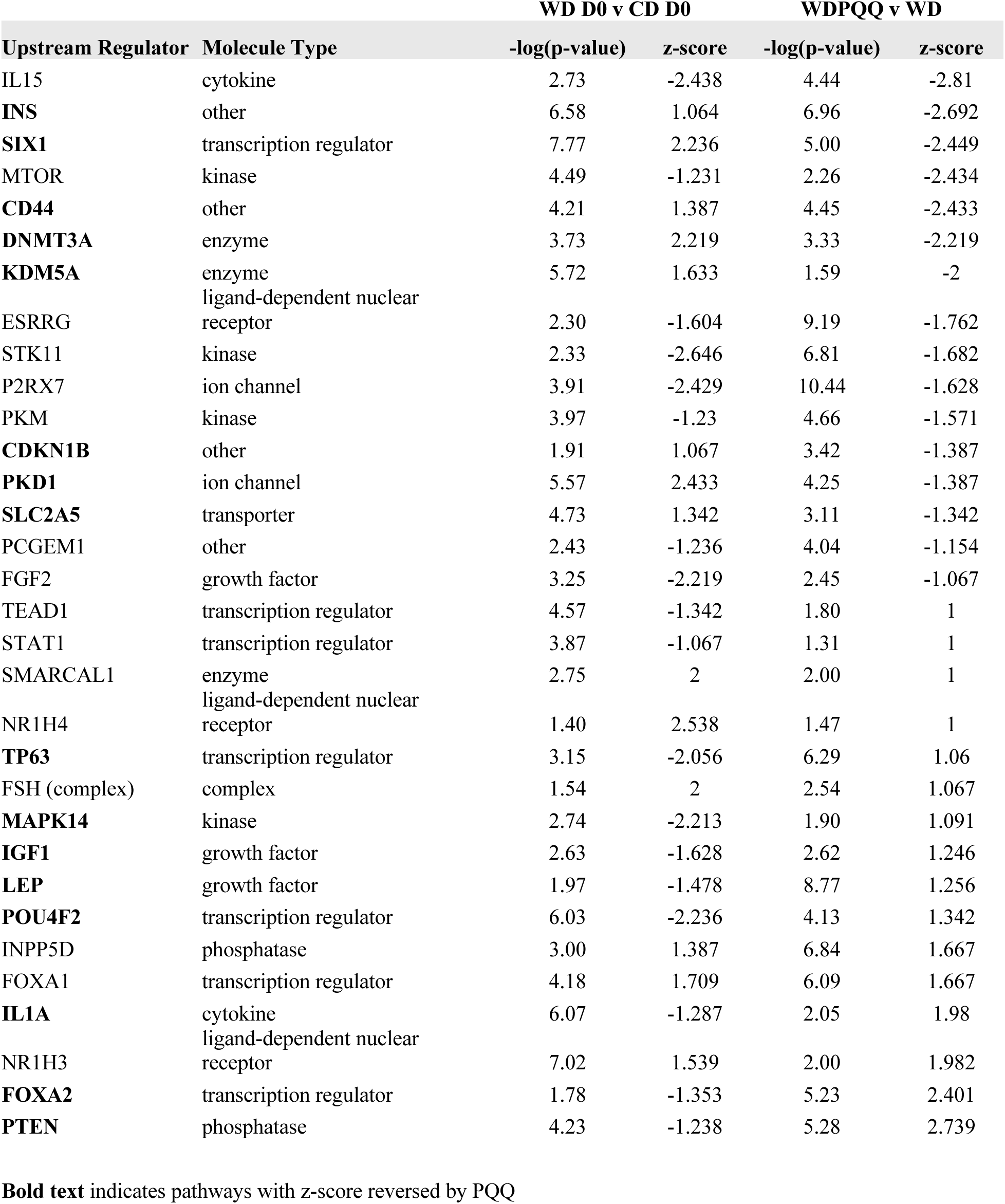
Enriched upstream regulators in plasma and serum.

We also investigated upstream regulator changes using a Comparison analysis between D0 and D30 (**Figure 4c**). PQQ supplementation did not affect diet-induced changes in several upstream regulators in plasma, including NR1H4, INSR, IL6, GRN, and OGA. PQQ markedly activated PPARGC1A, a known target, and activated PKM, P2RX7, IL15, PRL, and TP63. Regulators of pathways that are predicted to be significantly inhibited by PQQ include SKIC2 and HNF4A. These findings suggest that PQQ supplementation induced a diet-independent plasma signature of increased oxidative metabolism and remodeling of hepatic lipid metabolism associated with HNF4A signaling.

In WD D0 v CD D0 serum, using the same filters as in plasma, only two upstream regulators were identified (SLC13A1 and MYOD1), both with activation z-scores of 2. Therefore, stringency was reduced to −log10 p-value cutoff (corresponding to p<0.03) and |z-score| of 1.5 (**Figure 5a**). Using reduced stringency, the top enriched upstream regulator was IGF1R. Other upstream regulators of significantly activated pathways included HIF1A, CHUK, FHL2, MAP3K14, and NR1H4. Regulators of significantly inhibited pathways included PTEN, HUWE1, and FOXO1. At D30 (**Figure 5b**), PQQ supplementation to WD females resulted in activation of six pathways, including pathways regulated by NTRK1, CDKN1A, and ADIPOQ. Eight upstream regulators showed inhibition. Almost 30% of upstream regulators altered by diet or PQQ were transcriptional regulators, and 20% were kinases (**Supplementary Table 8**), suggesting PQQ targets pathways that may be epigenetically regulated. Comparison analysis between D0 and D30 (**Figure 5c**) showed more modest effects of PQQ in serum, except for a notable increase in PTEN activation and inhibition of RUNX1, suggesting PQQ may promote a shift toward reduced AKT signaling and inflammatory macrophage activation.

Representations of upstream regulator networks for FOXA2 (**Supplementary Fig. 4**) and NTRK1 (**Supplementary Fig. 5**) show that both FOXA2- and NTRK1-regulated signaling networks are predicted to be inhibited in WD compared with CD at D0 but are rescued by PQQ. These changes are linked with predicted PQQ-mediated increases in LXR signaling, IGF transport, and lipoprotein assembly, remodeling, and clearance. Signaling through xenobiotic transcription regulators PXR and AHR, PTEN, and TGF-beta receptor is predicted to be inhibited, as is signaling through IL-13. In contrast, IL-10 receptor antagonist-mediated signaling is activated in WD and inhibited by PQQ, decreasing TGFB1 and remodeling of the extracellular matrix (**Supplementary Fig. 6**). Together, these network-level changes suggest that PQQ restores FOXA2- and NTRK1-dependent metabolic signaling while suppressing pro-fibrotic, xenobiotic, and inflammatory pathways to restore homeostasis.

## DISCUSSION

In this study, we evaluated the effects of short-term (30 days) daily oral PQQ supplementation in non-pregnant adult female olive baboons (*Papio anubis*) on systemic metabolic and inflammatory markers. PQQ administration did not change body weights or adiposity assessed by sum of skin folds. We previously found that pregnant dams fed WD showed elevated triglycerides and cholesterol relative to pregnant CD-fed dams at 0.6 and 0.9 gestation. In contrast, non-pregnant animals in the present study did not exhibit significant baseline differences between diet groups in cholesterol (HDL-C, LDL-C, total cholesterol) or triglycerides, although HDL-C trended higher (p<0.1) in WD dams prior to PQQ supplementation. Mean triglycerides were also higher in the WD dams at baseline; however, high variability prevented statistical significance. PQQ supplementation for 30 days was effective in lowering serum HDL-C and LDL-C (p<0.05) and showed a trend for reducing total cholesterol (p<0.1) in WD dams. In CD dams, PQQ significantly lowered LDL-C. Triglyceride levels were not significantly altered in either group by PQQ. Based on these findings, once daily oral PQQ at a human-equivalent dose effectively lowered cholesterol in overweight/obese female primates consuming a high fat, high sugar, high cholesterol diet.

Obesity is well-recognized as a driver of chronic, low-grade systemic inflammation, through adipose tissue expansion, macrophage activation, and increased circulating inflammatory mediators ^29–31^. Serum cholesterol is a potent activator of monocytes, and in our study, we observed a significant effect of PQQ on both serum sCD163 and CRP. Because sCD163 is a marker for activated monocytes/macrophages, the PQQ-mediated reduction in serum sCD163 is consistent with our finding of lowered HDL-C and LDL-C in WD dams. PQQ also lowered CRP in both diet groups. Since CRP is primarily produced by the liver, this suggests that PQQ acts, at least in part, at the hepatic level to decrease inflammation. These results have important implications for PQQ and cardiometabolic health as they support the idea that PQQ use in overweight/obese women prior to pregnancy could yield substantial benefits for fetal health by improving maternal systemic metabolism and reducing inflammation.

Our proteomics, enrichment, and pathway analyses provide additional evidence that PQQ improves metabolic health by attenuating inflammatory signaling and modulating key metabolic pathways. At baseline, DAVID and STRING analyses showed WD-fed baboons exhibited marked upregulation and enrichment of inflammatory mediators, including the complement system and platelet activation pathways. After 30 days of PQQ supplementation, these same pathways were significantly downregulated, indicating a broad PQQ-mediated suppression of systemic inflammation in WD-fed animals. Notably, plasma levels of ankyrin repeat domain 2 (ANKRD2) – a regulator of muscle stress responses that promotes resolution of inflammatory signaling through inhibition of NF-κB signaling - were significantly reduced in WD animals at D0 but increased following PQQ supplementation.^32^. In addition, D30 serum from WD+PQQ animals showed upregulation of pathways involved in lipid metabolism, including *HDL and LDL remodeling, assembly, and clearance*; *chylomicron remodeling and assembly*; *assembly of lipase complexes*; *insulin like growth factor (IGFs) transport and uptake*; and *insulin signaling*. Together, these changes suggest potential mechanisms by which PQQ enhances lipid transport and clearance, attenuates inflammation, and improves insulin sensitivity, thereby countering negative immunometabolic effects of a high fat, high sugar diet.

To gain deeper insight into regulatory relationships beyond the enrichment patterns identified by DAVID and STRING, we performed IPA analyses, which highlighted key upstream regulators and canonical pathways modulated by PQQ in WD-fed animals. In the WD state, compared with CD, IPA revealed activation of upstream regulators such as SIX1 and KLF6, both linked to lipogenesis and fibrosis. SIX1 promotes *de novo* lipogenesis and liver fibrosis through TGF-β signaling and is implicated in MASLD progression, while KLF6 contributes to lipid biosynthetic pathways such as mTOR ^33^. WD baboons also showed inhibition of metabolic regulators including PPARGC1 and PTEN. Notably, 30 days of PQQ supplementation reversed these patterns. Activation of PPARGC1, a known PQQ target and central regulator of mitochondrial biogenesis and energy metabolism, together with downregulation of pro-lipogenic regulators like SIX1 and KLF6, suggests PQQ targets and promotes energy production through lipid metabolism while suppressing lipogenesis.

Thirty days of PQQ supplementation in WD-fed animals also led to predicted activation of pathways regulated by NRTK1, FOXA2 and ADIPOQ; these upstream regulators and associated pathways may represent potential novel targets of PQQ. Neurotrophic tyrosine kinase receptor 1 (NRTK1) encodes tyrosine kinase receptor A (TrkA), the high-affinity receptor for neurotrophin nerve growth factor (NGF). NGF-TrkA signaling is crucial for nerve growth and survival and is essential development of sympathetic neurons ^34^ ^35^. NGF is also an adipokine, produced and secreted by both white and brown adipose tissue ^36^. Within adipose tissue, NGF supports the growth and density of sympathetic nerve endings that innervate adipose tissue, forming neuro-adipose nexuses (NANs) that link adipocytes with these nerve endings. Stimulating NANs induces lipolysis and reduces fat mass, directly affecting lipid metabolism and energy homeostasis^37^ ^38^. NGF also exerts pro-inflammatory effects within adipose depots, likely via signaling through the low-affinity receptor p75^NTR^ ^38^. Notably, PQQ was previously shown to potently enhance NGF production *in vitro*, supporting a potential mechanistic link ^39^. Adiponectin, encoded by the ADIPOQ gene, is another key adipokine with potent anti-inflammatory, insulin-sensitizing and lipid regulating functions that are typically downregulated in obese and insulin resistant states ^40^. Interestingly, adiponectin contributes to the anti-depressant effects of exercise by promoting neurogenesis and neuroplasticity through brain derived neurotrophic factor (BDNF) signaling, a neurotrophin pathway similar to NTRK that signals through the TrkB receptor ^41^. Together, these findings suggest that PQQ may modulate neurotrophin- and adipokine-mediated pathways that integrate metabolic, inflammatory, and neural regulation of energy homeostasis.

Another important upstream regulator predicted to be activated by PQQ supplementation in WD-fed animals is Forkhead Box Protein A2 (FOXA2; also known as Hepatic Nuclear Factor 3 Beta: HNF3β), a member of the FOXA family of transcription factors, required for the normal development and function of liver and pancreas, key metabolic organs ^42^. FOXA2 controls genes responsible for cellular metabolism by regulating glucose and lipid metabolism and plays a key role in adipocyte differentiation ^43,44^. FOXA2 deficiency in hepatocytes causes hepatic lipid accumulation, decreased glucose uptake, glycogen storage, and insulin resistance. FOXA2 is downregulated in metabolic conditions like MASLD. FOXA2 is a "pioneer" transcription factor that opens chromatin at bile acid and drug-metabolism loci and cooperates with or is antagonized by xenobiotic nuclear receptors, conjugating enzymes, and transporters ^45–47^. FOXA2 acts as a master regulator for hepatic metabolism and inflammation. Its dysfunction is a central mechanism connecting insulin resistance, dyslipidemia, and chronic inflammation. In metabolic disease, insulin-driven Akt signaling represses FOXA2 ^45^ ^48^ and nuclear lamina remodeling redistributes FOXA2 binding ^49^, permanently altering the liver’s epigenetic landscape to favor lipid storage, reduced beta oxidation, and chronic inflammation. Notably, FOXA2 has been identified in the human fetus circulating proteome ^50^. FOXA2 normally activates genes such as Cpt1a, Hmgcs2 ^48^, and Mtp ^51^, supports HDL formation through ApoM ^52^ and VLDL secretion ^51^, and directly suppresses TLR4 and NF- κ B–driven cytokine production ^53,54^; loss of FOXA2 therefore promotes dyslipidemia and chronic inflammation.

As a pioneer factor, FOXA2 maintains chromatin accessibility for other transcription factors regulating bile acid transporters, detoxification genes, and drug-metabolizing enzymes ^46^. FOXA2 also interacts with the Pregnane X Receptor (PXR), a ligand-activated nuclear receptor traditionally known for detoxifying xenobiotics but increasingly recognized as a critical regulator of glucose, lipid, and immune homeostasis ^55^. Ligand-activated PXR represses FOXA2s oxidative programs ^55^; moreover, chromatin accessibility, receptor occupancy, and ligand responsive gene activation by xenobiotic nuclear receptors such as FXR and LXR- α require FOXA2, with FOXA2 occupancy markedly increasing upon agonist treatment and forming ligand-dependent complexes with these receptors ^56^. Through these reciprocal actions, FOXA2 integrates nutrient, inflammatory, and xenobiotic cues to coordinate hepatic metabolic homeostasis.

The upstream regulators activated by PQQ may converge on the sympathetic nervous system (SNS), which plays a central role in maintaining energy homeostasis through changing energy states and physiological demands. Depending on physiological stimuli, SNS activation in the cardiovascular system, liver, pancreas, skeletal muscle, and adipose tissue regulates blood pressure, resting metabolic rate, thermogenesis, and both glucose and lipid metabolism. In the WD-fed state, PQQ may mitigate early signs of metabolic dysfunction by promoting production of mature NGF and enhancing NGF-TrkA signaling rather than pro-NGF-p75^TRK^, supporting sympathetic innervation of adipose tissue and improved lipid and glucose handling, potentially through downstream metabolic mediators such as FOXA2 (**Figure 6**). Consistent with this model, WD+PQQ animals showed reduced levels of inflammatory markers, suggesting PQQ may also engage anti-inflammatory pathways, including those mediated by adiponectin. Although NGF is often elevated in the obese state due to TNF-α-driven inflammation ^57,58^, NGF-mediated sympathetic activity can promote beneficial metabolic effects, including stimulating production of leptin and adiponectin, increasing thermogenesis, browning of white fat, lipolysis, and increasing expression of PPARγ, a known target of PQQ ^34,38,59,60^. Thus, PQQ’s ability to reverse WD-induced metabolic stress, lower CRP and sCD163, and improve lipid profiles may reflect activation of NGF/SNS circuits within adipose tissue. Future studies directly testing the connection between NGF signaling, peripheral SNS activity, and PQQ treatment will be essential to evaluate this hypothesis.

**Figure 6.**
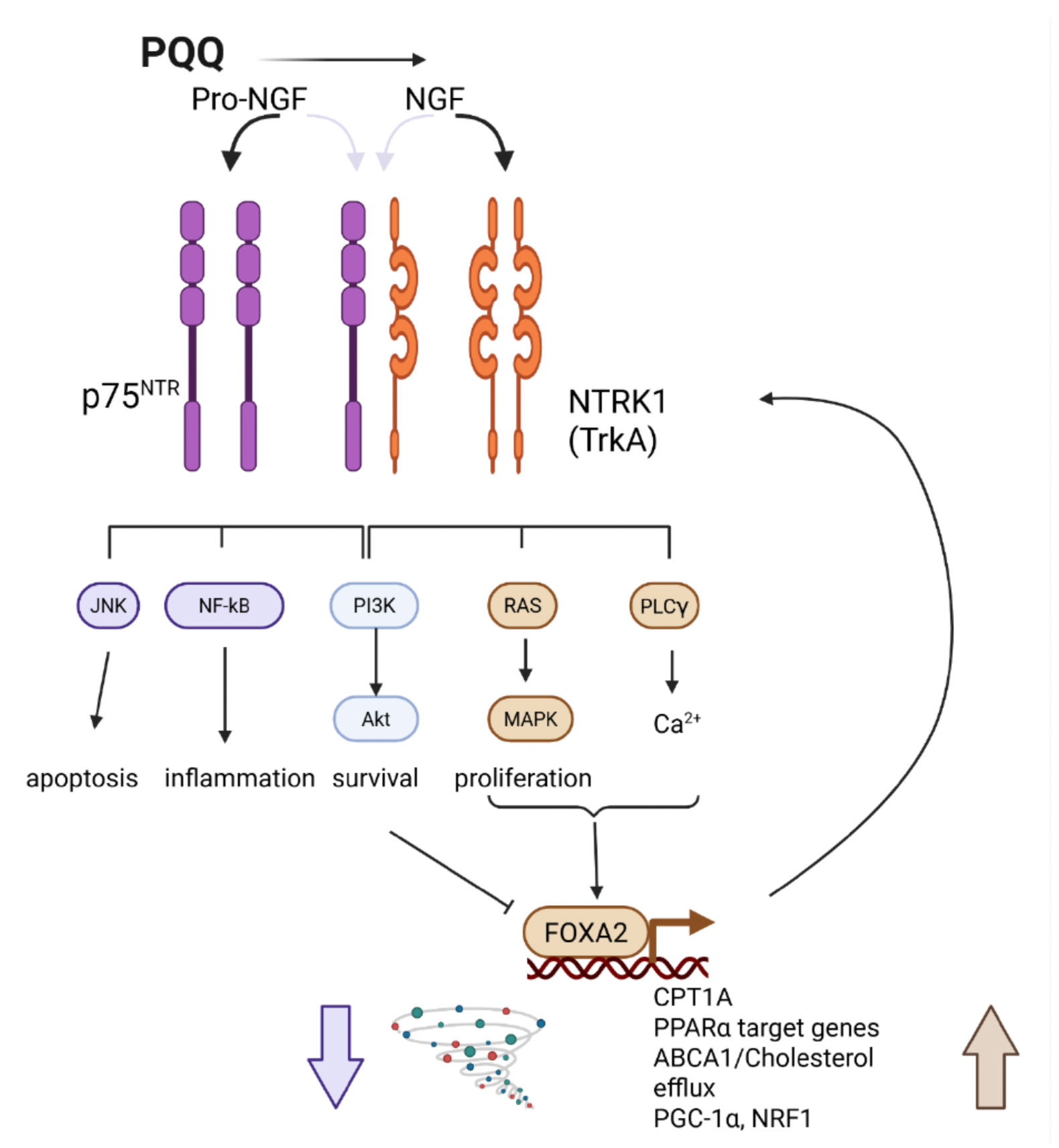
Proposed model by which PQQ promotes activation of signaling through NTRK1 and FOXA2 to reduce inflammation and improve lipid metabolism. PQQ enhances production of NGF. We postulate that PQQ drives signaling through NTRK1 rather than through p75NTR, reducing inflammation and cytokines. Downstream pathways associated with proliferation, as well as calcium signaling, activate FOXA2 to promote increased transcription of genes involved in lipid metabolism and transport (CPT1A, PPARα, ABCA1) as well as genes promoting mitochondrial function (PGC-1α, NRF1). FOXA2 itself activates NTRK1 signaling, which is modulated when NTRK1 forms a heterodimer with p75^NTR^ to activate PI3K and Akt signaling and inhibit FOXA2. Adapted from Samario-Roman, et al (PMID: 36768281). Created with Biorender.

A major strength of this study is the use of a well-characterized NHP model that closely recapitulates human metabolic physiology, dietary responses, and immune regulation, enhancing translational relevance. The paired, within-animal design and integration of clinical metabolic markers with unbiased plasma and serum proteomics, followed by complementary pathway analyses (DAVID, STRING, and IPA), provide a unique systems-level view of PQQ’s immunometabolic effects. In addition, the use of a human-equivalent PQQ dose supports clinical relevance. However, several limitations should be acknowledged. The small cohort size, inherent to NHP studies, limits statistical power and necessitated relaxed thresholds for some proteomic analyses. The 30-day intervention period captures early metabolic and inflammatory remodeling but may be insufficient to assess longer-term effects on adiposity, insulin sensitivity, or tissue-specific outcomes. Moreover, conclusions regarding mechanistic pathways, including sympathetic nervous system involvement and FOXA2- or NTRK1-mediated signaling, are based on predictive bioinformatic analyses rather than direct functional measurements in relevant tissues. Future studies incorporating larger cohorts, longer treatment durations, and direct assessment of tissue-specific signaling and sympathetic activity will be essential to validate and extend these findings.

## MATERIALS AND METHODS

### Animals and Ethical Approval

All experiments adhered to the guidelines as outlined by the U.S. National Institutes of Health Office of Laboratory Animal Welfare Public Health Service Policy on Humane Care and Use of Laboratory Animals and the Animal Welfare Act for the housing and care of laboratory animals. The research was conducted with approval from the University of Oklahoma Health Sciences Center Institutional Animal Care and Use Committee (IACUC; 25-024-SAF). Environmental enrichment was provided for all animals.

Adult female baboons (*Papio anubis*) were maintained in two separate social groups based on diet for the study’s duration (**Figure 1a and 1b**). One cohort (Control Diet; CD; n=5, 8-9 years of age) were fed a standard monkey chow (5045*, Purina LabDiets, St. Louis, MO; 30.4% calories from protein, 13.3% calories from fat, 56.4% calories from carbohydrates; metabolizable energy: 3.14 kcal/g). The second cohort (n=4; 7-9 years of age) were maintained on a Western-style Diet (WD; TAD Primate diet, 5L0P; Purina LabDiets, St. Louis, MO; 18% calories from protein, 36.3% calories from fat, 45.4% from calories from carbohydrates; metabolizable energy: 3.40kcal/g). The WD-fed females were provided access to a high-fructose (100g/L) beverage (Kool-Aid^TM)^ *ad libitum*. Both diets were otherwise matched in micro and macro-nutritional content. The WD cohort had been maintained on the WD/fructose drink diet for approximately 3 years. All dams received equal daily dietary enrichment (e.g., fruit, peanuts). Four of the CD females were multiparous (2 prior pregnancies; but were not presently nursing) and one CD female was nulliparous. Three of the WD females were multiparous (n=2 with two prior pregnancies; n=1 with one prior pregnancy; but were not presently nursing) while one WD female was nulliparous.

Blood (plasma and serum) and anthropometric measurements (body weight, crown length, skin fold measurements [thigh, triceps, subscapular, abdominal] and girth) were obtained under Ketamine sedation prior to and after the 30 day-PQQ administration period. PQQ (BioPQQ®, a gift from Mitsubishi Gas and Chemical) was administered once daily to each female at 0.25 mg/kg body weight (allometrically scaled to the recommended human dose of 20 mg/day). PQQ dose was based on body weight measured at the start of the study. PQQ in powdered form was added to a highly palatable “cake ball” treat. The female baboons in this study were trained by the research staff to take the treats by hand. Each female was observed to eat the treat, ensuring accurate dosing of each animal.

### Serum C-Reactive Protein (CRP), soluble CD163 (sCD163), Triglyceride and Cholesterol ELISAs

CRP and soluble CD163 were measured using a C-Reactive Protein human serum ELISA kit (EIA-3954, DRG International, INC, Springfield, NJ) and a human CD163 ELISA kit (DC1630, R&D Systems, Minneapolis, MN), respectively. Briefly, a 1:100 (CRP) and 1:10 (sCD163) serum dilution of Day 0 (pre-PQQ) and D30 samples was performed in duplicate and added to the antibody pre-coated plates available in the kits. The assays were performed using the manufacturer’s instructions. The concentration of CRP and sCD163 in the serum samples was determined by reading absorbance at 450 nm.

Serum triglycerides were measured in duplicate with 1:2 serum dilution using a Triglyceride colorimetric assay (10010303, Cayman Chemical, Ann Arbor, MI). This assay measures triglyceride concentration in serum by enzymatic hydrolysis of triglycerides to glycerol and free fatty acids. Colorimetric measurement of the glycerol released was obtained by measuring absorbance at 540nm.

Serum cholesterol was measured using EnzyChrom^TM^ HDL and LDL/VLDL Assay Kit (EHDL-100, BioAssay Systems, Hayward, CA). The samples and cholesterol standard (300mg/dL) were prepared according to the manufacturer’s instructions. 20 ul of undiluted serum for the HDL and LDL/VLDL fractions and 12 ul of undiluted serum for the total cholesterol fraction per animal were used for the assay. The final concentration of the cholesterol standard and samples were measured by reading absorbance at 340nm.

### Proteomics

#### S-Trap Methods (Serum/Plasma) – Orbitrap Eclipse DIA

Proteomics on D0 and D30 plasma and serum samples were performed at the University of Arkansas for the Medical Sciences (UAMS) Institutional Development Award (IDeA) National Resource for Quantitative Proteomics (Little Rock, AR). Following depletion of abundant proteins from plasma or serum samples using High Select Top 14 resin (Thermo), remaining proteins were reduced, alkylated, and digested with sequencing grade modified porcine trypsin (Promega) using S-Trap 96-well plates following the protocol from the manufacturer (ProtiFi). Tryptic peptides were then separated by reverse phase XSelect CSH C18 2.5 um resin (Waters) on an in-line 150 x 0.075 mm column using an UltiMate 3000 RSLCnano system (Thermo). Peptides were eluted using a 60 min gradient from 98:2 to 65:35 buffer A:B ratio. Eluted peptides were ionized by electrospray (2.2 kV) followed by mass spectrometric analysis on an Orbitrap Eclipse Tribrid mass spectrometer (Thermo). To assemble a chromatogram library, six gas-phase fractions were acquired on the Orbitrap Eclipse with 4 m/z DIA spectra (4 m/z precursor isolation windows at 30,000 resolution, normalized AGC target 100%, maximum inject time 66 ms) using a staggered window pattern from narrow mass ranges using optimized window placements. Precursor spectra were acquired after each DIA duty cycle, spanning the m/z range of the gas-phase fraction (i.e., 496-602 m/z, 60,000 resolution, normalized AGC target 100%, maximum injection time 50 ms). For wide-window acquisitions, the Orbitrap Eclipse was configured to acquire a precursor scan (385-1015 m/z, 60,000 resolution, normalized AGC target 100%, maximum injection time 50 ms) followed by 50x 12 m/z DIA spectra (12 m/z precursor isolation windows at 15,000 resolution, normalized AGC target 100%, maximum injection time 33 ms) using a staggered window pattern with optimized window placements. Precursor spectra were acquired after each DIA duty cycle. Buffer A = 0.1% formic acid, 0.5% acetonitrile, Buffer B = 0.1% formic acid, 99.9% acetonitrile

#### Data Analysis for Proteomics

Following data acquisition, data were searched using Spectronaut (Biognosys version 18.3) against the UniProt *Papio anubis* database (Proteome ID: UP000028761, 4^th^ version of 2023) using the directDIA method with an identification precursor and protein q-value cutoff of 1%, generate decoys set to true, the protein inference workflow set to maxLFQ, inference algorithm set to IDPicker, quantity level set to MS2, cross-run normalization set to false, and the protein grouping quantification set to median peptide and precursor quantity. Protein MS2 intensity values were assessed for quality using ProteiNorm ^61^. The data was normalized using Cyclic Loess ^62^ and analyzed using proteoDA to perform statistical analysis using Linear Models for Microarray Data (limma) with empirical Bayes (eBayes) smoothing to the standard errors ^62^. Proteins with an FDR adjusted p-value < 0.05 and a fold change > 2 were considered significant. Expression of ten proteins in the comparison between CDPQQ v CD and two proteins in the comparison between WD v CD were significantly different in plasma (Table S1), and there were no significant differences between groups in serum. Therefore, for the purpose of carrying out pathway analysis, significance was relaxed and a raw p < 0.1 and log2 fold change > 1.5 was used for subsequent analyses. UniProt IDs were mapped to gene symbols. IDs that were not mapped were searched using BLAST restricted to Human taxonomy. The top scoring entry was selected. Duplicates were removed, and entries were selected based on the p value for pairwise comparisons or the Spectronaut score. The Statistical Analysis [One-Factor] module of MetaboAnalyst 6.0 (https://www.metaboanalyst.ca) was used for PLS-DA analyses, t-tests, and hierarchical clustering ^63^. Features with more than 50% missing values were excluded, and the Interquantile range filter for variance was set to 10%. Limit of Detection (LoD) data estimation was used for zero-filling. Data were Log2 transformed and mean centered to normalize.

Meta-analysis of the functional annotation clustering of the proteomics data was done using the Database for Annotation, Visualization and Integrated Discovery or DAVID ^64,65^ and the STRING database of predicted functional associations between proteins ^66^. Annotations for analysis using DAVID include UP_KW_PTM, GOTERM_BP_DIRECT, GOTERM_CC_DIRECT, GOTERM_MR_DIRECT, and INTERPRO. Functional Annotation Clustering was performed separately on upregulated and downregulated proteins for each comparison in plasma and serum. Options selected were Kappa Similarity Term Overlap set to 3, Similarity Threshold of 0.3, Initial and Final Group Membership restricted to 2, multiple linkage threshold of 0.3, and ease set to 0.5. If gene names could not be mapped using Official Gene Symbol, conversion to Entrez gene ID was performed, using Papio Anubis as the reference background. Finalized tables include entries with FDR < 0.05. Duplicate entries with the same genes within a cluster were deleted. Upregulated and downregulated proteins were also analyzed separately with STRING. Cutoffs used were p<0.1 and absolute value of fold change > 0.5. Proteins were added to the network until PPI Enrichment <1×10^-16^. Up to 15 Reactome pathways were plotted based on “strength”, and pathways with similarity > 0.5 were merged when more than 15 pathways were identified with significant FDR.

Canonical pathway and upstream regulator analysis of the proteomics data was carried out using Qiagen’s Ingenuity Pathway Analysis (IPA) (Qiagen, Redwood City, CA). The number of features used in each analysis following data reduction are summarized in **Supplementary Table 1**.

## STATISTICAL ANALYSIS

Serum CRP, sCD163, TGs, HDL, LDL/VLDL cholesterol, and the sum of skin folds were analyzed by ANOVA with Tukey’s post-hoc analysis. Body weights were analyzed by Student’s t-test. Significance was set at p ≤ 0.05.

## DATA AVAILABILITY

The mass spectrometry proteomics data generated in this study have been deposited to the ProteomeXchange Consortium via the PRIDE partner repository with the dataset identifier PXD072309 and 10.6019/PXD072309. All other data generated in this study are provided in the Supplementary Information.

## CODE AVAILABILITY

This study did not generate any unique code.

## ACKNOWLEDGEMENTS

IDeA National Resource for Quantitative Proteomics and NIH/NIGMS grant R24GM137786.802 Presbyterian Health Foundation Bridge Grant (KRJ) and NIH/NIDDK R01 DK139443 (KRJ and DAM).

## AUTHOR CONTRIBUTIONS

SGD, DAM and KRJ wrote, reviewed, and edited the manuscript and contributed to data analysis and interpretation. DAM and KRJ designed all experiments. KRJ analyzed and visualized proteomics data provided by the University of Arkansas for the Medical Sciences (UAMS) Institutional Development Award (IDeA) National Resource for Quantitative Proteomics Facility. SGD and KEH performed and analyzed immunological and metabolic assays. JFP supervised all animal work. DNR, TLS, and ABT provided PQQ supplementation and collected samples and anthropometric measurements.

