## Supplementary figures and images for "Thirty days of supplementation with PQQ reprograms immunometabolic networks in Western diet-fed female baboons"

### Supplemental Figure S1

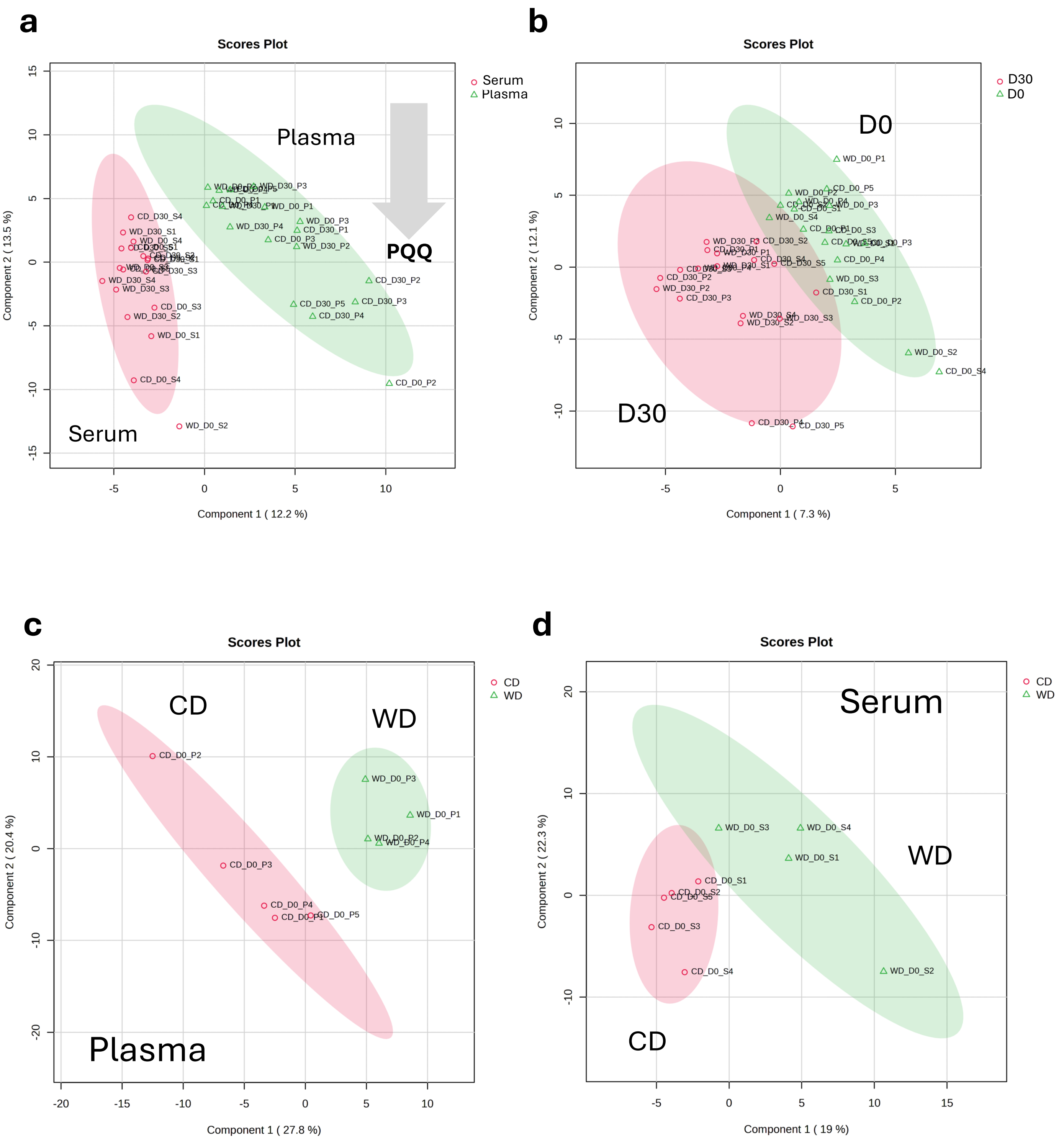


Supplementary Figure S1

### Supplemental Figure S2

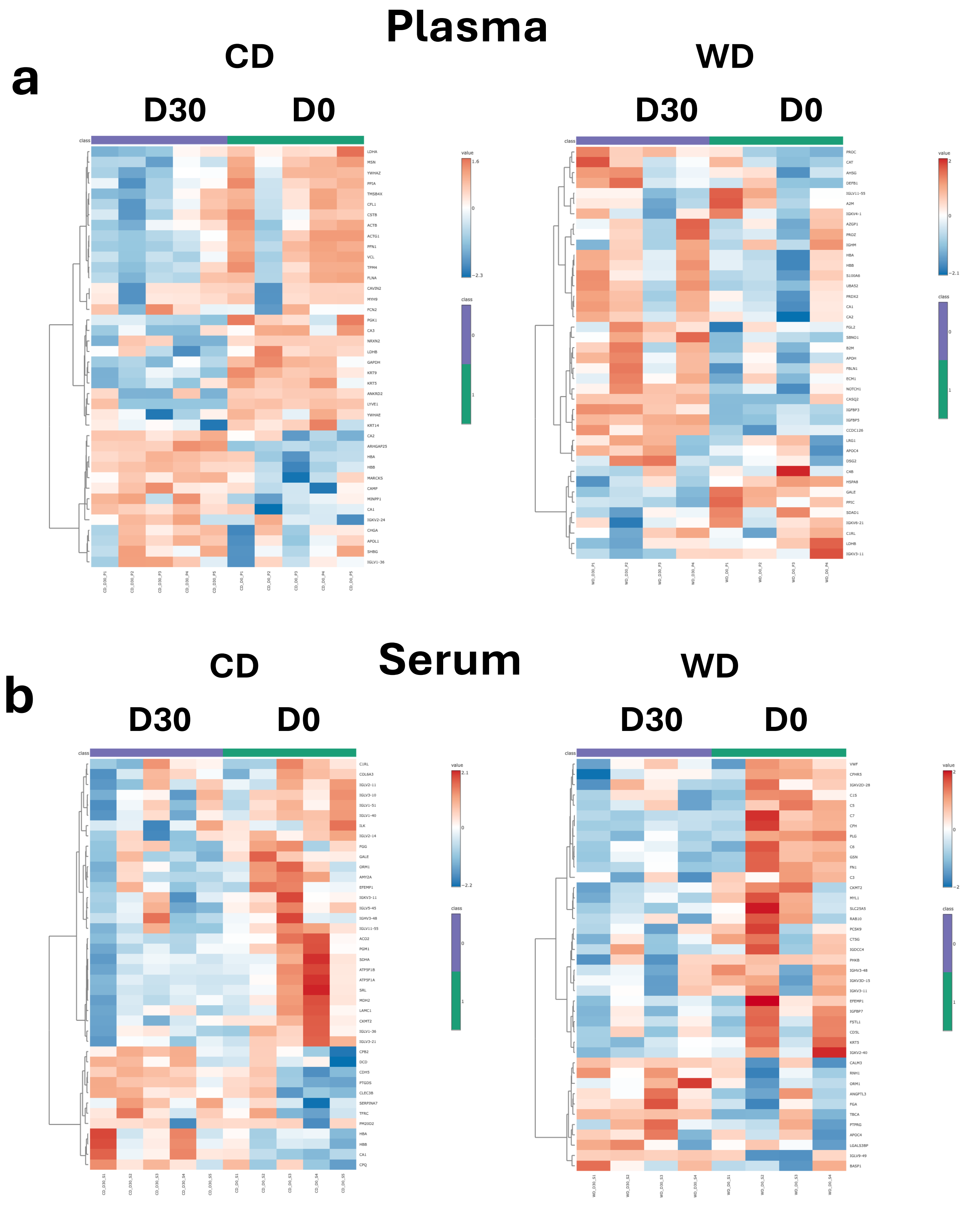


Supplementary Figure S2

### Supplemental Figure S3

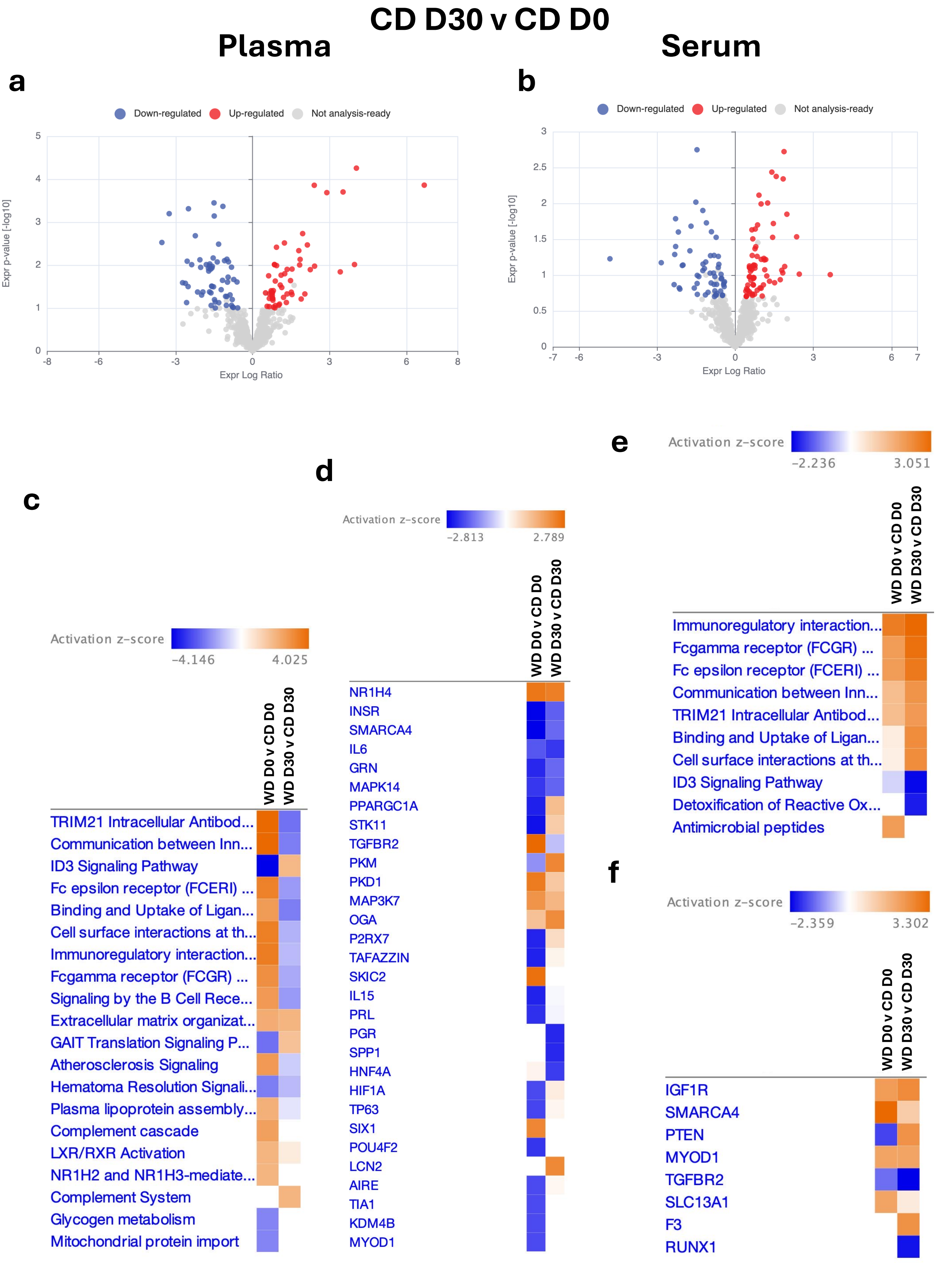


Supplementary Figure S3

### Supplemental Figure S4

**a**

WD D0 v CD D0

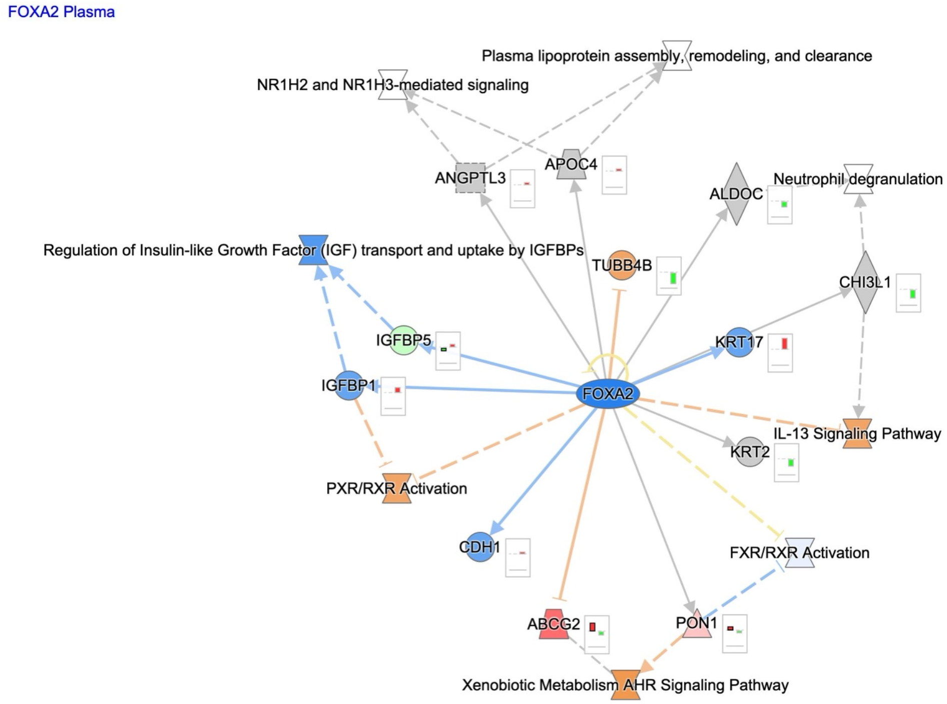


**b**

WD D30 v WD D0

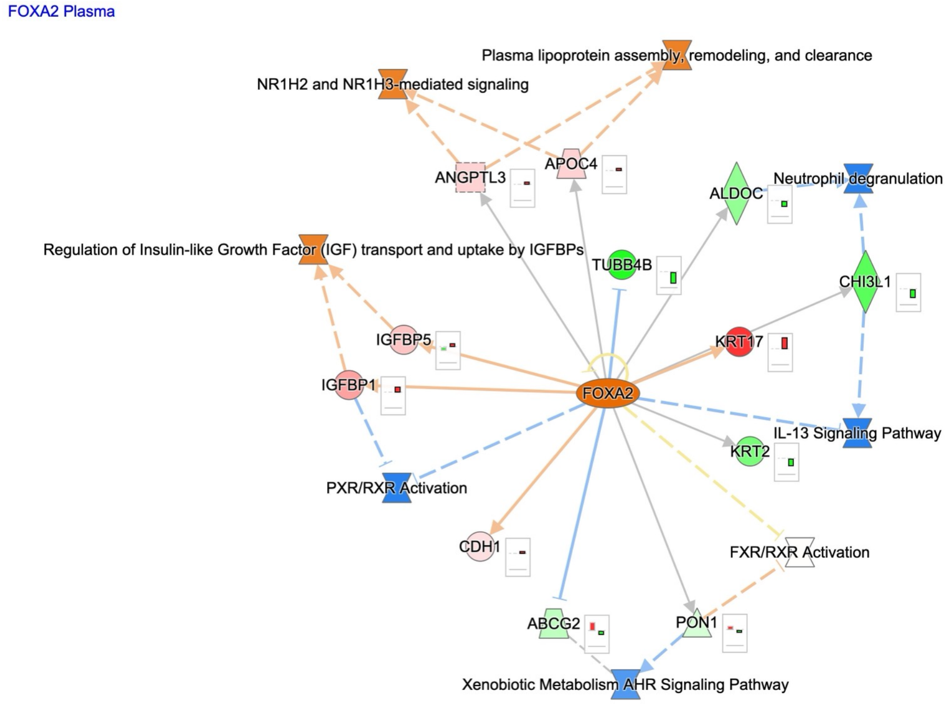


Supplementary Figure S4

### Supplemental Figure S5

**a**

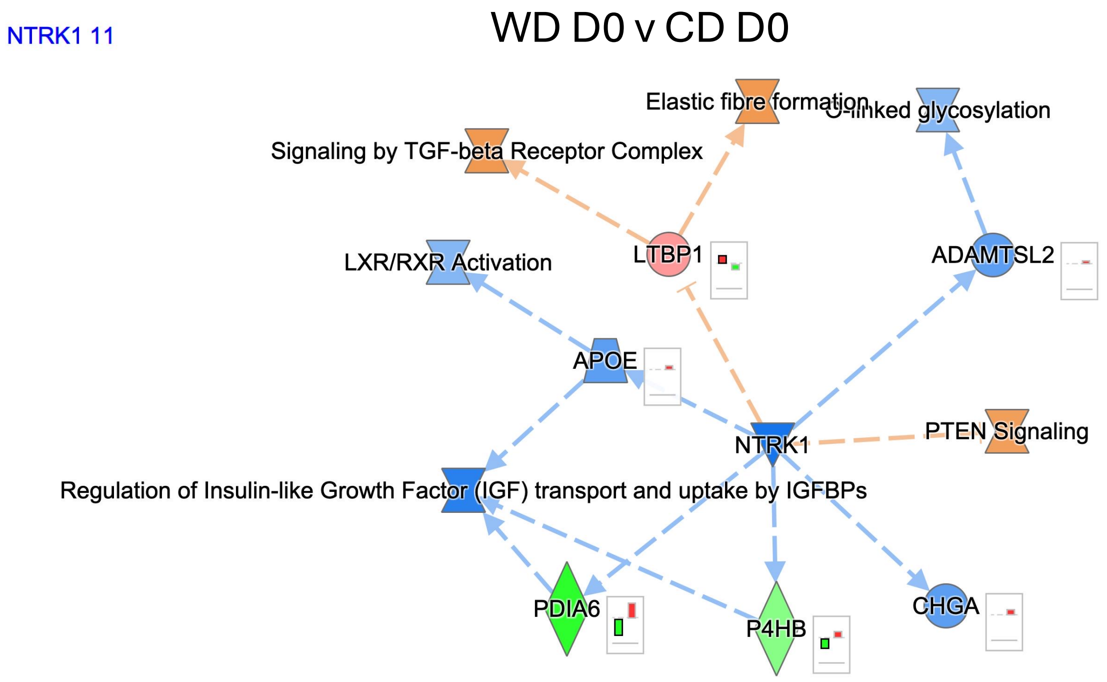


**b**

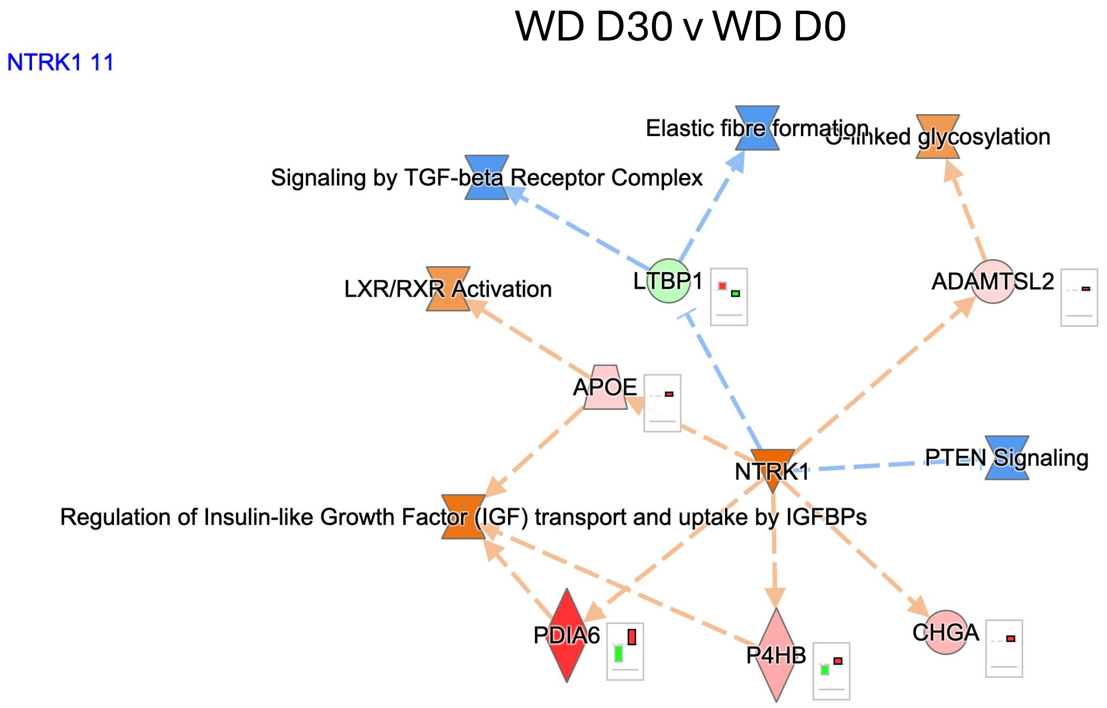


Supplementary Figure S5

### Supplemental Figure S6

**a**

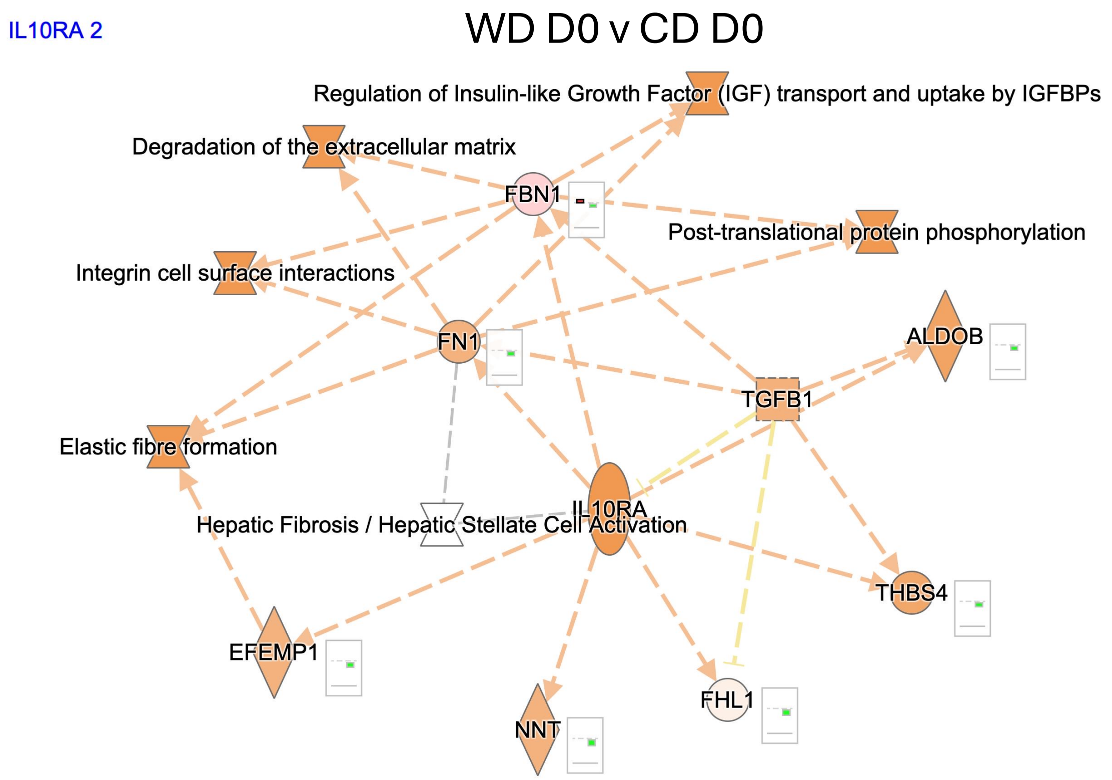


**b**

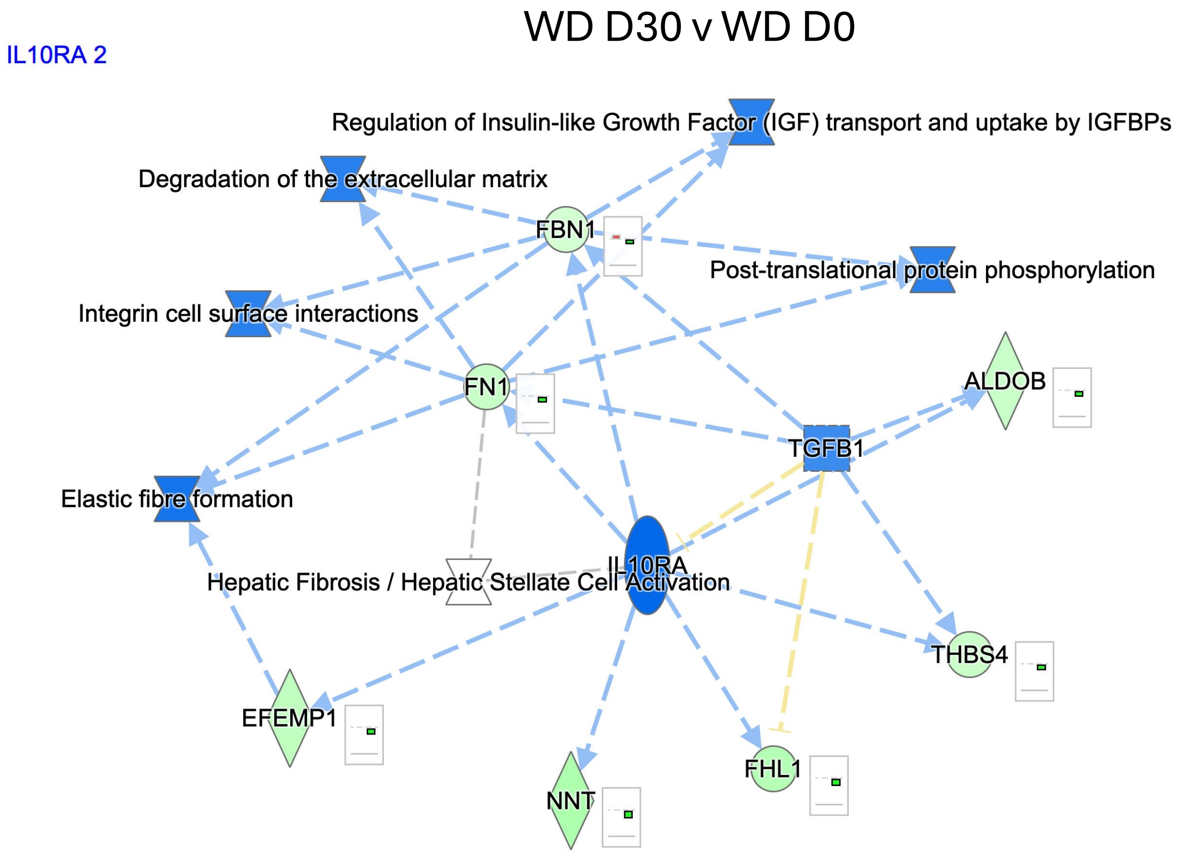


Supplementary Figure S6
